# Calnexin mediates the maturation of GPI-anchors through ER retention

**DOI:** 10.1101/2020.07.08.192955

**Authors:** Xin-Yu Guo, Yi-Shi Liu, Xiao-Dong Gao, Taroh Kinoshita, Morihisa Fujita

**Affiliations:** Key Laboratory of Carbohydrate Chemistry and Biotechnology, Ministry of Education, School of Biotechnology, Jiangnan University, Wuxi, Jiangsu 214122, China; Research Institute for Microbial Diseases, Osaka University, Suita, Osaka 565-0871, Japan; WPI Immunology Frontier Research Center, Osaka University, Suita, Osaka 565-0871, Japan

## Abstract

The protein folding and lipid moiety status of glycosylphosphatidylinositol-anchored proteins (GPI-APs) are monitored in the endoplasmic reticulum (ER), with calnexin playing dual roles in the maturation of GPI-APs. In the present study, we investigated the functions of calnexin in the quality control and lipid remodeling of GPI-APs in the ER. By directly binding the N-glycan on proteins, calnexin was observed to efficiently retain GPI-APs in the ER until they were correctly folded. In addition, sufficient ER retention time was crucial for GPI-inositol deacylation, which is mediated by post-GPI attachment protein 1 (PGAP1). Once the calnexin/calreticulin cycle was disrupted, misfolded and inositol-acylated GPI-APs could not be retained in the ER and were exposed on the plasma membrane. In calnexin/calreticulin deficient cells, endogenous GPI-anchored alkaline phosphatase was expressed on the cell surface, but its activity was significantly decreased. ER stress induced surface expression of misfolded GPI-APs, but proper GPI-inositol deacylation occurred due to the extended time that they were retained in the ER. Our results indicate that calnexin-mediated ER quality control systems for GPI-APs are necessary for both protein folding and GPI-inositol deacylation.

## Introduction

Glycosylphosphatidylinositol (GPI) is a complex glycolipid that acts as a membrane anchor for many cell surface proteins (Ikezawa, 2002; McConville & Menon, 2000; Tiede, Bastisch, Schubert, Orlean, & Schmidt, 1999) and is a highly conserved post-translational modification from yeast to mammals. In mammals, there are at least 150 GPI-anchored proteins (GPI-APs), which serve as receptors, adhesion molecules, enzymes, protease inhibitors, and so on (Kinoshita, 2020). The biosynthesis of GPI and its attachment to protein occur in the endoplasmic reticulum (ER). Nascent GPI-APs synthesized by GPI transamidase are still immature and undergo remodeling reactions to become mature GPI-APs. In the ER, two remodeling reactions occur in many GPI-APs. First, an acyl-chain on an inositol-ring of the GPI-anchor is eliminated by the GPI-inositol deacylase PGAP1 (Chen et al., 1998; Tanaka, Maeda, Tashima, & Kinoshita, 2004). Second, a side-chain ethanolamine-phosphate (EtNP) attached on the second mannose (Man2) of GPI-glycan is removed by the GPI-EtNP phosphodiesterase PGAP5 (Fujita et al., 2009). These remodeling reactions are crucial for the interaction of GPI-APs with p24 protein complexes, which are cargo receptors for GPI-APs (Fujita & Kinoshita, 2012; Fujita et al., 2011), indicating that GPI-AP remodeling in the ER is required for their efficient sorting into transport vesicles at the ER exit sites. Pathogenic homozygous mutations in *PGAP1* cause an inherited GPI deficiency, which results in intellectual disability, encephalopathy and hypotonia (Granzow et al., 2015; Kettwig et al., 2016; Murakami et al., 2014). It has been reported that *Pgap1* mutant mice exhibit abnormal head development, such as otocephaly (Ueda et al., 2007; Zoltewicz et al., 2009) and holoprosencephaly (McKean & Niswander, 2012), suggesting that Pgap1 function is required for normal forebrain formation. In addition, male *Pgap1*-knockout mice are infertile (Ueda et al., 2007). Taken together, these findings show that correct processing of GPI-anchors is crucial for the proper functions of these proteins in vivo.

Protein asparagine (N)-glycosylation is another type of post-translational modification occurring in the ER (Hammond, Braakman, & Helenius, 1994; Schwarz & Aebi, 2011). An oligosaccharide consisting of Glc_3_Man_9_GlcNAc_2_ (Glc, glucose; Man, mannose, GlcNAc, *N*-acetylglucosamine) is transferred to the amino group of asparagine within the motif NxS/T (N, asparagine; S/T, serine or threonine; x, any amino acids except proline) of newly synthesized proteins. The folding states of secretory proteins are monitored by the ER quality control systems (Sun & Brodsky, 2019). N-glycans are processed in the ER, which contributes to protein folding. First, two Glc residues on N-glycans are trimmed by α-glucosidase I and II. Processed monoglucosylated N-glycan structures are then recognized by calnexin and calreticulin, which are molecular chaperones that possess a lectin domain. Calnexin has a transmembrane domain, whereas calreticulin is a soluble protein (Helenius & Aebi, 2004), and both proteins associate with protein disulfide isomerases such as ERp57 and ERp29, prompting the folding of newly synthesized proteins (Ellgaard & Frickel, 2003; Kozlov, Muñoz-Escobar, Castro, & Gehring, 2017). Once the remaining Glc residues on protein N-glycans are trimmed by α-glucosidase II, calnexin/calreticulin dissociate from the proteins. However, proteins remaining in an unfolded state are re-glucosylated by UDP-Glc: glycoprotein glucosyltransferase (UGGT), subsequently becoming bound again to calnexin and calreticulin for refolding. This series of reactions is called the calnexin/calreticulin cycle and is essential for glycoprotein folding.

In our previous study, we performed a genetic screening to identify factors that affect GPI-inositol deacylation. In the screening, *MOGS*, *GANAB*, and *CANX*, as well as *PGAP1*, were identified as candidates (Liu et al., 2018). *MOGS*, *GANAB*, and *CANX* encode α-glucosidase I and II and calnexin, respectively (Tannous, Pisoni, Hebert, & Molinari, 2015). These results suggested that N-glycan-dependent ER quality control systems participate in the lipid remodeling of GPI-APs, whereas it was unclear how calnexin contributes to the processing.

When proteins fail to fold correctly, they are recognized as misfolded proteins, the majority of which are retained in the ER and degraded through the ER-associated degradation (ERAD) pathway (Vembar & Brodsky, 2008). However, misfolded GPI-APs do not appear to be suitable substrates for ERAD, possibly because of the presence of GPI-anchors. A fraction of misfolded GPI-APs are degraded through proteasomes (Fujita, Yoko-o, & Jigami, 2006; Ishida et al., 2003), whereas the rest are exported from the ER and delivered to the vacuole for degradation in yeast (Hirayama, Fujita, Yoko-o, & Jigami, 2008; Sikorska et al., 2016). In mammalian cells, although misfolded GPI-APs are retained in the ER, they are rapidly released into the secretory pathway upon acute ER stress despite their misfolding (Satpute-Krishnan et al., 2014). Time-lapse imaging of live cells under acute ER stress conditions showed misfolded prion proteins are transported from the ER to the Golgi and plasma membrane, and subsequently to lysosomes for degradation.

In the present study, we investigated the roles of calnexin in the quality control and lipid remodeling of GPI-APs in the ER. In wild-type (WT) cells, GPI-APs were folded and inositol deacylated in the ER and expressed at the cell surface. In calnexin and calreticulin double knockout (CANX&CALR-DKO) cells, protein folding and GPI-inositol deacylation efficiencies were significantly decreased such that misfolded and inositol-acylated GPI-APs were exposed on the plasma membrane. Thus, these results indicate that the N-glycan-dependent calnexin/calreticulin cycle is responsible for the correct folding of GPI-APs and provides sufficient ER retention time for efficient GPI-inositol deacylation.

## Materials and Methods

### Cells, antibodies, and materials

All the cell lines, antibodies, and other reagents are listed in the key resource table. HEK293 and HEK293FF6 cells (Hirata et al., 2015) and their knockout (KO) derivatives were cultured in Dulbecco’s Modified Eagle medium (DMEM) containing 10% (vol/vol) FCS (Biological Industries). G418 (400 μg/ml), puromycin (1 μg/ml), and streptomycin/penicillin (100 units/mL of penicillin and 100 μg/mL of streptomycin) were used where necessary. Cells were maintained at 37°C under a humidified atmosphere with 5% CO_2_. CANX&CALR-DKO cells stably expressing CANX WT or mutants were selected with 10 μg/ml blasticidin.

Mouse monoclonal anti-CD59 (clone 5H8), anti-CD55 (clone IA10) (Liu et al., 2018; Maeda et al., 2007), anti-FLAG (M2; Sigma), anti-RGS-His6 (BSA-free; QIAGEN), anti-calnexin (M178-3; MBL), and anti-GAPDH (clone 1E6D9; Proteintech) antibodies were used as primary antibodies. Other primary antibodies used in the present study included rabbit monoclonal anti-calreticulin (clone D3E6; Cell Signaling Technology), polyclonal anti-calnexin (C4731; Sigma), polyclonal anti-ERp57 (Proteintech), polyclonal anti-GRP78 (Novus), polyclonal anti-eGFP (Proteintech), monoclonal anti-ALPP (clone SP15; Abcam) and anti-HA-Tag (C29F4; Cell Signaling Technology). Phycoerythrin (PE)-conjugated goat anti-mouse IgG (eBioscience), PE-conjugated donkey anti-rabbit IgG (eBioscience), HRP-conjugated anti-mouse IgG (Cell Signaling Technology) and anti-rabbit IgG (Cell Signaling Technology), F(ab’)2-Goat anti-Mouse IgG (H+L) Cross-Adsorbed Secondary Antibody, Alexa Fluor 555 (Thermo Fisher Scientific), and F(ab’)2-Goat anti-Mouse IgG (H+L) Cross-Adsorbed Secondary Antibody, Alexa Fluor 488 (Thermo Fisher Scientific) were used as secondary antibodies. PE-conjugated anti-CD109 (HU17; eBioscience) was used for flow cytometry. Thapsigargin (100 nM, TG, Sigma-Aldrich) and doxycycline (1 μg/ml, 631311; Clontech Laboratories) were used for drug treatments.

### Plasmids

All the plasmids and oligonucleotides used in this study are listed in Table S1 and Table S2, respectively. For the CRISPR-Cas9 system used to KO target genes, guide RNA sequences were designed using the E-CRISP website (Heigwer, Kerr, & Boutros, 2014) (http://www.e-crisp.org/E-CRISP/), and the corresponding DNA fragments were ligated into the *Bpi*I-digested vector pX330-EGFP. The CANX cDNA fragment was amplified from human cDNA and cloned into the retroviral vector pLIB2-BSD to generate pLIB2-BSD-CANX. The lectin-deficient mutant plasmids pLIB2-BSD-CANX (Y164A), pLIB2-BSD-CANX (K166A), pLIB2-BSD-CANX (M188A) and pLIB2-BSD-CANX (E216A) and the PDI interaction mutant plasmids pLIB2-BSD-CANX (D343A), pLIB2-BSD-CANX (E351A), pLIB2-BSD-CANX (D343A, E351A), pLIB2-BSD-CANX (W342A, D343A), and pLIB2-BSD-CANX (D347A, E351A) were constructed by site-direct mutagenesis. The DNA fragments encoding mEGFP-FLAG-CD59 and mEGFP-FLAG-CD55 were digested with *Eco*RI and *Not*I from pPB-FRT-PGKp-BSD-mEGFP-FLAG-CD59 and pPB-FRT-PGKp-BSD-mEGFP-FLAG-CD55 (Liu et al., 2018), and then cloned into pME-BSD-mEGFP-FLAG-CD59 and pME-BSD-mEGFP-FLAG-CD55 respectively. The misfolded, soluble, and N-glycan-deficient mutants of CD59 were constructed by site-directed mutagenesis. The transmembrane portion of CD46 was amplified from CD46 cDNA and ligated to pME-BSD-EGFP-FLAG-BSD-CD59-(C94S) to generate CD59-(C94S)-TM. For localization, pME-Zeo-mRFP-KDEL (Fujita et al., 2009) was used as an ER marker. The LY6D and CD52 cDNA fragment was amplified from human cDNA and cloned into the retroviral vector pLIB2-His6-IRES2-BFP to generate pLIB2-His6-LY6D-IRES2-BFP and pLIB2-His6-CD52-IRES2-BFP. The plasmid pME-VSVG^ts^-FF-mEGFP-GPI (VFG-GPI) encoded a reporter protein consisting of the extracellular domain of a temperature-sensitive vesicular stomatitis virus G (VSVG^ts^) protein, a furin cleavage site, a FLAG tag, mEGFP (modified EGFP) and a GPI attachment signal (Fujita et al., 2009; Fujita et al., 2011; Maeda, Ide, Koike, Uchiyama, & Kinoshita, 2008; Takida, Maeda, & Kinoshita, 2008).

### Establishment of knockout cell lines

To generate *PDIA3* gene knockout cell lines, HEK293 cells were transiently transfected with two pX330-PDIA3 plasmids (cr1 and cr2) (Table S1) bearing different gRNAs targeting to exon regions of *PDIA3* gene. After 3 days, cells with EGFP were sorted using a cell sorter S3e (Bio-Rad). Then, the collected cells were cultured for 8 days and subjected to limiting dilution and obtain the clonal KO cells. Clones lacking wild-type alleles of the target gene were selected, and DNA sequences were analyzed using the Sanger method.

### PI-PLC treatment and flow cytometry analysis

For phosphatidylinositol-specific phospholipase C (PI-PLC; Thermo Fisher Scientific) treatment of endogenous GPI-APs, 10^6^ cells were harvested with trypsin/EDTA. Then, the cells were incubated with 5 U/ml of bacterial PI-PLC dissolved in PBS supplemented with 0.5% BSA, 5 mM EDTA, and 10 mM HEPES (pH 7.4) plus DMEM medium without FCS for 1.5 h at 37°C. After being washed with PBS, the cells were stained with the specific primary antibodies for endogenous CD59, CD55 and CD109 (10 μg/ml) for 25 min on ice and then washed two times with cold FACS buffer (PBS containing 1% BSA and 0.1% NaN_3_). Subsequently, the cells were incubated with PE-conjugated goat anti-mouse IgG as the secondary antibody for 25 min on ice, washed two times with FACS buffer and then analyzed with an Accuri C6 instrument (BD). The data were analyzed using FlowJo, and the remaining GPI-APs were calculated from mean values of PE-fluorescent intensity treated with or without PI-PLC.

For exogenous GPI-APs, pME-HA-GPI-AP (Table S1) was transfected together with pME-BFP, which acts as a transfection control, into WT and CANX&CALR-DKO cells. A retroviral plasmid vector containing His6-tagged GPI-AP-IRES2-BFP sequence was transfected into HEK293T packaging cells, and subsequently infected into WT and CANX&CALR-DKO cells. Lipofectamine 2000 (Thermo Fischer Scientific) was used as the transfection reagent. Three days after transfection, 10^6^ cells were harvested and treated with or without PI-PLC, and anti-HA or anti-His6 was used as the primary antibody. Subsequently, secondary antibody staining and flow cytometry analysis were performed as described above.

### Immunofluorescence analysis

To detect the subcellular localization of the EGFP-FLAG-tagged misfolded CD59 (EGFP-FLAG-CD59 (C94S)), N-glycan-deficient mutant CD59 (EGFP-FLAG-CD59 (C94S, N43Q)), transmembrane misfolded CD59 (EGFP-FLAG-CD59 (C94S) TM), misfolded CD55 (EGFP-FLAG-CD55 (C81A)), and N-glycan mutant CD55 (EGFP-FLAG-CD55 (C81A, N95Q)), HEK293FF6 and its KO derivative cells were transfected with plasmids expressing these proteins together with the plasmid pME-mRFP-KDEL as an ER marker. Then, 36 h after transfection, cells were replated onto glass coverslips pretreated with 1% gelatin and cultured for another 2 days. Subsequently, the cells were washed with PBS, fixed in 4% paraformaldehyde for 10 min at roon temperature, washed again with PBS, and then incubated with 40 mM ammonium chloride for 10 min at room temperature. Finally, the coverslips were mounted onto slides using a mounting solution containing DAPI for 5 min.

To assess the localization of VFG-GPI, HEK293FF6, CANX&CALR-DKO, and CANX&CALR-DKO+CANX cells were transfected with pME-mRFP-KDEL. Subsequently, 36 h after transfection, the cells were harvested and replated onto glass coverslips pretreated with 1% gelatin at 37°C for 1 day and then incubated with 1 μg/ml doxycycline at 40°C for 24 h to induce VFG-GPI expression. Then, the cells were quickly fixed with 4% paraformaldehyde for 10 min before being washed with PBS and then 40 mM ammonium chloride.

To assess the localization of endogenous calnexin, the cells were fixed and permeabilized with −20°C methanol for 5 min at 4°C before being blocked with PBS containing 5% FCS (blocking buffer) for 1 h. Subsequently, the cells were incubated for 1 h with mouse-anti-calnexin (M178-3; MBL) as primary antibody diluted in blocking buffer. Subsequently, the cells were gently washed with PBS twice. Then the cells were incubated for 1 h with an F(ab’)2-Goat anti-Mouse IgG (H+L) cross-adsorbed secondary antibody, Alexa Fluor 555 (Invitrogen, Thermo Fisher Scientific) diluted in blocking buffer, after which the cells were gently washed with PBS twice.

To assess the localization of HLA1-1-147-FLAG, pME-HLA1-1-147-FLAG and pME-mRFP-KDEL were transfected together into WT cells. Then, 36 h after transfection, the cells were harvested and replated onto glass coverslips pretreated with 1% gelatin and incubated at 37°C for 1 day followed by an incubation at 37°C with 100 nM TG for 6 h. The cells were then fixed with 4% paraformaldehyde, washed with PBS, and incubated with 40 mM ammonium chloride for 10 min at room temperature. Subsequently, the cells were permeabilized with 0.2% Triton-X 100 diluted in blocking buffer for 15 min at room temperature. Mouse-anti-FLAG (M2; Sigma-Aldrich) was used as the primary antibody and was diluted in blocking buffer for 1 h, after which the cells were gently washed with PBS twice. Then, the cells were incubated for 1 h with an F(ab’)2-Goat anti-Mouse IgG (H+L) cross-adsorbed secondary antibody conjugated with Alexa Fluor 488 (Invitrogen, Thermo Fisher Scientific) diluted in blocking buffer. Subsequently, the cells were gently washed with PBS twice.

The cells were visualized using a confocal microscope (C2si; Nikon) with a CFI Plan Apochromat VC oil objective lens (100× magnification and 1.4 NA). Pearson’s correlation coefficient (GFP/RFP) was analyzed by ImageJ using the JACoP plug-in.

### Immunoprecipitation

HEK293 cells cultured in 10-cm dishes were transfected with 16 μg of plasmids using Lipofectamine 2000 and incubated for 48 h. Then, the cells were harvested with trypsin/EDTA, washed with cold PBS at 4°C, and then incubated with 550 μl of lysis buffer (25 mM HEPES pH 7.4, 150 mM NaCl, 0.5% CHAPS, protein inhibitor cocktail (EDTA-Free, MCE) and 1 mM PMSF) on ice for 30 min. After incubation, the tube was centrifuged at 10,000 ×*g* for 10 min at 4°C to remove insoluble fractions. Then, a portion of the supernatant was mixed with sample buffer, the rest was transferred to a new tube, and then 20 μl of prewashed anti-FLAG affinity gel (A2220; Sigma-Aldrich) was added. After rotating the tubes at 4°C for 1-1.5 h, the gel with tagged proteins were washed 4 times with lysis buffer. The proteins were eluted by boiling in SDS sample buffer and analyzed by Western blotting.

### Cell surface alkaline phosphatase (ALP) activity assay

For both HEK293FF6 WT and CANX&CALR-DKO cells, 8,000 cells/well were seeded in 96-well plates and cultured overnight at 37°C. After the medium was removed, the cells were gently washed with 1×HBSS buffer (Ca^2+^ and Mg^2+^ free, pH 7.4; Sangon Biotech) twice, after which 150 μl/well of a pNPP solution (Alkaline Phosphatase Yellow (pNPP) Liquid Substrate System for ELISA, P7998; Sigma) was added. Then, the plates were incubated at 37°C for 6.5 h, the reaction was stopped by adding 50 μl/well of 3 M NaOH, and the absorbance at 405 nm was measured using an Enspire 2300 instrument. The calf intestinal alkaline phosphatase (TaKaRa) was used as a standard for the reaction.

## Results

### Calnexin/calreticulin cycle is required for efficient GPI-inositol deacylation for both endogenous and exogenous N-glycosylated GPI-APs

After GPI attachment to proteins, an acyl-chain linked to inositol on many GPI-APs is removed by the GPI-inositol deacylase PGAP1. The inositol-deacylation status of GPI-APs can be determined by assessing their sensitivity to bacterial PI-specific phospholipase C (PI-PLC) (Ferguson, Kinoshita, & Hart, 2009; Kmenon, 1994). PI-PLC reacts with the hydroxyl group on the 2-position of the inositol ring, producing inositol-1,2-cyclic phosphate (Heinz, Ryan, Bullock, & Griffith, 1995). If an acyl-chain is attached to the 2-position of the inositol, PI-PLC cannot cleave the substrate (Figure 1A). We previously demonstrated that GPI-inositol deacylation of CD59 and CD55 is partially impaired in cells defective for calnexin (CANX) and is more strongly affected in cells defective for both CANX and calreticulin (CALR) (Liu et al., 2018). To determine if the CANX/CALR-dependency of GPI-inositol diacylation is a general phenomenon of various GPI-APs, we treated CANX&CALR-DKO HEK293 cells with PI-PLC, and endogenous CD59, CD55, and CD109 in the CANX&CALR-DKO cells showed PI-PLC resistance (Figure 1B and C). To confirm whether the PI-PLC resistance of GPI-APs is a general phenomenon, exogenous human GPI-APs CD48, CD14, ART3, ART4, LY6E, LY6G6C, and CD52 were transiently expressed and assessed by PI-PLC treatment. Consistent with the endogenous GPI-APs, the exogenous GPI-APs also showed PI-PLC resistance in CANX&CALR-DKO cells (Figure 1D and Figure 1-figure supplement). These results suggest that disruption of the calnexin/calreticulin cycle causes inefficient GPI-inositol deacylation in GPI-APs.

**Figure 1.**
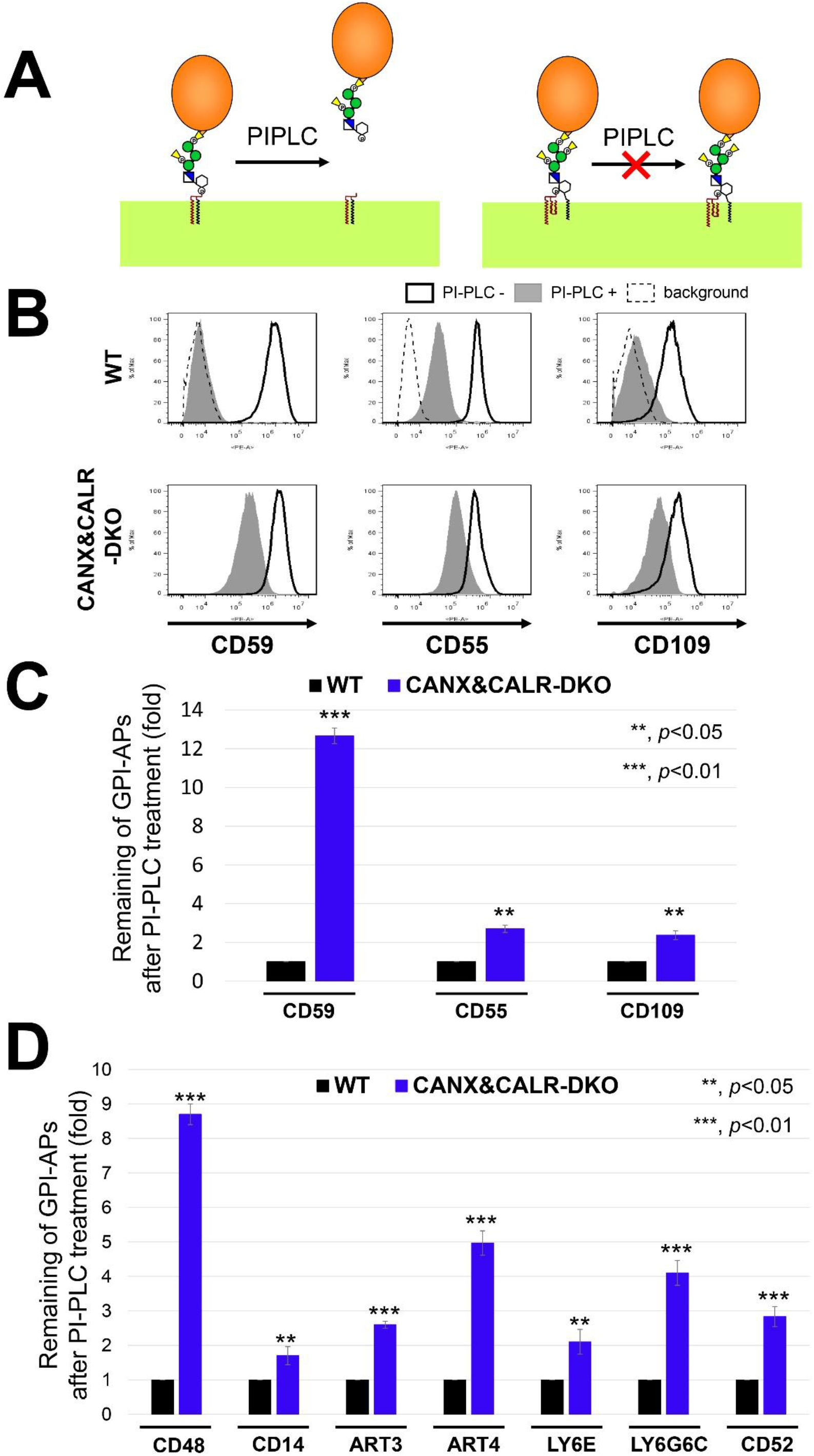
GPI-inositol-acylated GPI-APs are expressed on the plasma membrane in CANX&CALR-DKO cells. (A) After lipid remodeling, mature GPI-APs on the cell surface can be cleaved and released from the plasma membrane by PI-PLC (left). When an acyl-chain is modified to the 2-position of the inositol ring on GPI-APs, PI-PLC cannot cleave the GPI-APs (right). (B) WT and CANX&CALR-DKO cells were treated with or without PI-PLC. Endogenous CD59, CD55 and CD109 were stained with the appropriate antibodies and analyzed by flow cytometry. Gray shaded areas indicate cells treated with PI-PLC, heavy solid lines indicate cells without PI-PLC treatment, and dashed lines show the background of the secondary antibody. (C) Remaining endogenous GPI-APs after PI-PLC treatment. The fluorescence intensity of GPI-APs after PI-PLC treatment was divided by that observed without PI-PLC treatment. The values in WT cells were set as 1, and the relative values for CANX&CALR-DKO were plotted. The data are presented as the means ± SD of three independent measurements. *P*-values (one-tailed, student’s t-test) are shown. (D) Remaining exogenous GPI-APs after PI-PLC treatment. HA-tagged GPI-AP constructs were co-transfected with a BFP-expressing plasmid into WT and CANX&CALR-DKO cells. A retroviral vector containing His6-tagged CD52-IRES2-BFP was stably expressed in WT and CANX&CALR-DKO cells. The BFP-positive regions were gated, and cell surface expression of GPI-APs were analyzed by flow cytometry. The values were calculated as described in (C).

Approximately 95% of GPI-APs in mammalian cells contain at least one N-glycan (Liu et al., 2018). To determine whether calnexin/calreticulin also affects the inositol-deacylation of non-N-glycosylated GPI-APs, we analyzed LY6D and GFP-GPI, which have no N-glycan, by PI-PLC treatment in WT and CANX&CALR-DKO cells. In WT cells, both LY6D and GFP-GPI were sensitive to PI-PLC cleavage (Figure 2A, left top), whereas in CANX&CALR-DKO cells, unlike N-glycosylated GPI-APs, LY6D and GFP-GPI did not show PI-PLC resistance (Figure 2A, left bottom, right). We next constructed HA-tagged CD59 with (WT) or without N-glycan (N43Q) to compare the PI-PLC sensitivity between WT and CANX&CALR-DKO cells. HA-CD59 WT showed PI-PLC resistance in CANX&CALR-DKO cells, compared to that observed in WT cells (Figure 2B, left). The PI-PLC sensitivity of HA-CD59 (N43Q) was comparable between the WT and CANX&CALR-DKO cells (Figure 2B, right). The observed difference between N-glycosylated and non-N-glycosylated GPI-APs suggests that GPI-inositol deacylation regulation by calnexin/calreticulin is dependent on the N-glycans on GPI-APs.

**Figure 2.**
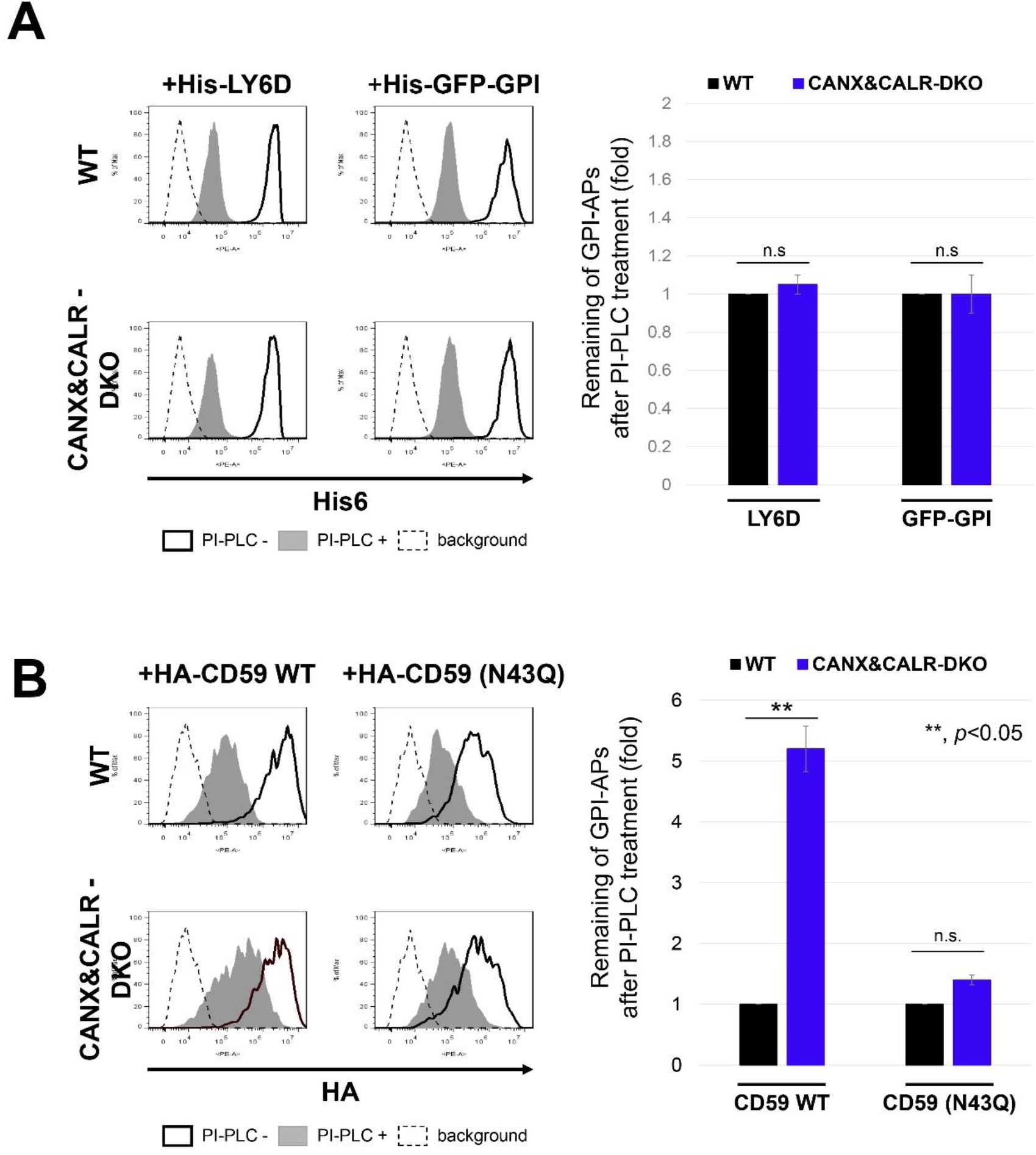
Non-N-glycosylated GPI-APs use a calnexin-independent mechanism for GPI-inositol deacylation. (A) The plasmids pLIB2-His6-LY6D-IRES-BFP or pLIB2-His6-GFP-GPI-IRES-BFP were packaged into retroviruses and infected into WT and CANX&CALR-DKO cells. Cells were treated with or without PI-PLC after 3 d of transfection. Anti-His6 was used as the primary antibody to detect the surface levels of His6-LY6D and His6-GFP-GPI. The cells showing the same BFP intensities were gated, and the surface expression was analyzed by flow cytometry (left). The values for the remaining GPI-APs after PI-PLC treatment in WT cells were set as 1, and the relative values in CANX&CALR-DKO cells were plotted (right). The data are presented as the means ± SD of three independent measurements. *P*-values (one-tailed, student’s t-test) are shown. (B) WT or non-N-glycosylated mutant constructs (pME-HA-CD59 and pME-HA-CD59 (N43Q), respectively) were transfected into cells together with pME-BFP (transfection control). Three days after transfection, the cells were treated with or without PI-PLC, then HA-tagged CD59 was analyzed by staining with an anti-HA antibody (left). Cells showing the same BFP intensities were gated for the same transfection level. The values for the remaining HA-CD59 WT or N43Q after PI-PLC treatment in WT cells were set as 1, and the relative values in CANX&CALR-DKO cells were plotted (right). The data are presented as the means ± SD of three independent measurements. *P*-values (one-tailed, student’s t-test) are shown.

### Glycan binding of calnexin is required for efficient GPI-inositol deacylation

The structure of the ER-luminal portion of calnexin has two domains: a glycan binding domain, consisting of a globular β-sandwich structure, and an extended arm domain (P-domain), consisting of two β-strands (Schrag et al., 2001). The glycan binding domain preferentially recognizes an N-glycan containing Glc_1_Man_9_GlcNAc_2_ (Spiro, Zhu, Bhoyroo, & Soling, 1996; Vassilakos, Michalak, Lehrman, & Williams, 1998). The amino acids Y164, K166, M188, and E216 (Michael R Leach & Williams, 2004) have been reported to be important for the lectin activity of calnexin. To assess whether this activity affects GPI-inositol deacylation, we constructed calnexin constructs with mutations in the amino acids responsible for glycan binding (Y164A, K166A, M188A, and E216A). Compared with a wild-type calnexin construct, the PI-PLC sensitivity of CD59 in CANX&CALR-DKO cells transfected with glycan binding-deficient calnexin constructs was rarely rescued, which was consistent with previous results (Y164A and E216A) (Liu et al., 2018) (Figure 3A and Figure 3-figure supplement A and B). The results indicated that lectin activity of calnexin is necessary for efficient GPI-inositol deacylation.

**Figure 3.**
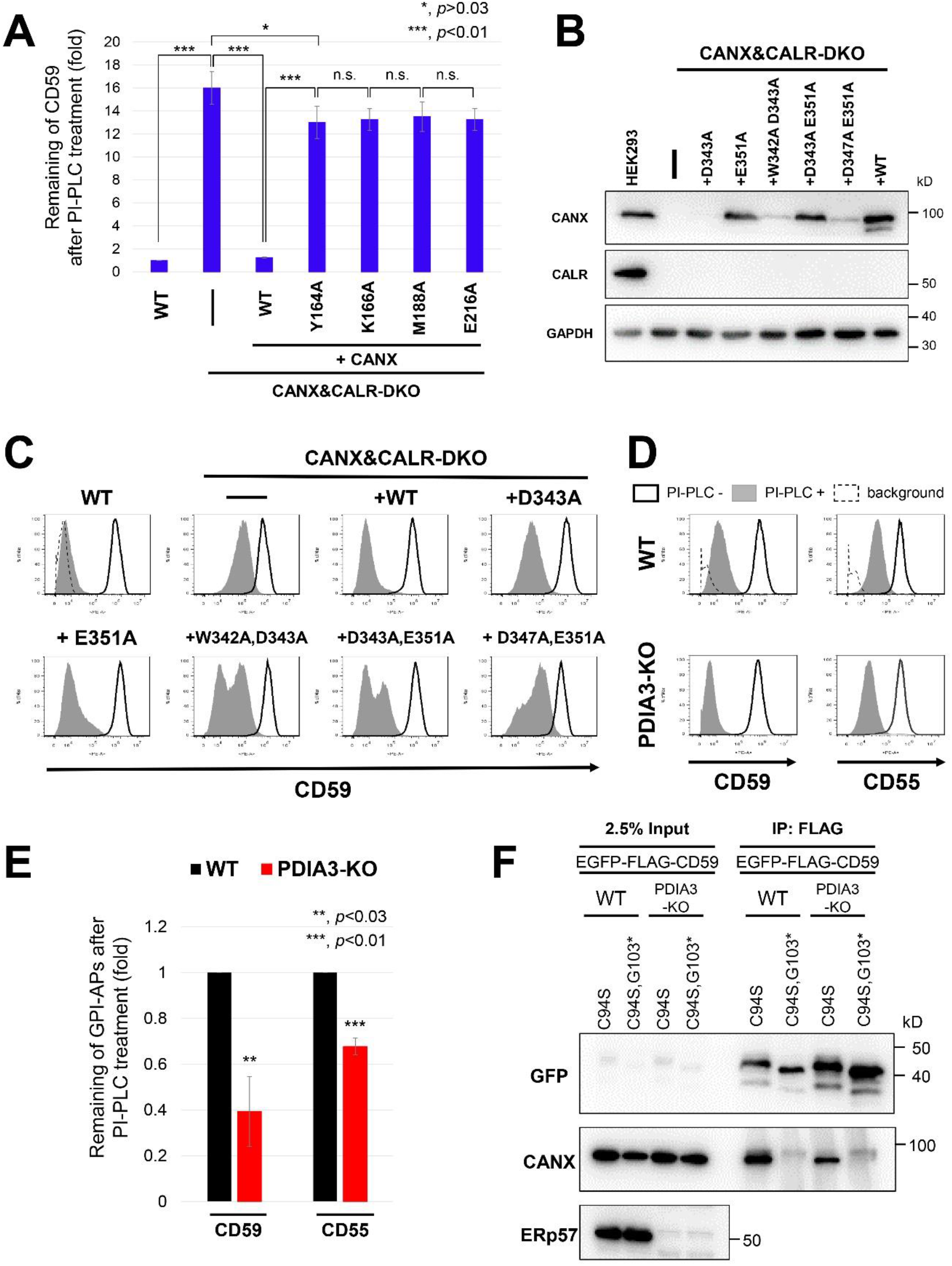
Lectin but not ERp57 binding of calnexin is required for efficient GPI-inositol deacylation. (A) Calnexin mutants defective in lectin activity did not rescue the PI-PLC resistance in CANX&CALR-DKO cells. WT and CANX&CALR-DKO cells stably expressing WT CANX or lectin activity-deficient CANX mutants (Y164A, K166A, M188A or E216A) were treated with or without PI-PLC. The relative intensities of remaining CD59 were plotted. The value in WT cells was set as 1. The data are presented as the means ± SD of three independent measurements. *P*-values (two-tailed, student’s t-test) are shown. (B) WT and CANX&CALR-DKO cells stably expressing WT CANX or CANX mutants defective in ERp57 binding (D343A, E351A, W342A&D343A, D343A&E351A or D347A&E351A) were lysed, after which calnexin and calreticulin were detected by immunoblotting. GAPDH was used as a loading control. (C) Calnexin mutants defective in ERp57 and ERp29 binding could rescue the PI-PLC resistance in CANX&CALR-DKO cells. WT and CANX&CALR-DKO cells stably expressing WT CANX or CANX mutants defective in ERp57 binding (D343A, E351A, W342A&D343A, D343A&E351A or D347A&E351A) were treated with or without PI-PLC. The surface expression of CD59 was analyzed by flow cytometry as described in Figure 1B. (D and E) GPI-APs in PDIA3-KO cells showed greater PI-PLC sensitivity than that observed for the parental WT cells. WT and PDIA3-KO cells were treated with or without PI-PLC. After treatment, the surface expression of CD59 and CD55 was analyzed (D) as described in Figure 1B. The levels of CD59 and CD55 remaining after PI-PLC treatment were plotted. The values for the remaining CD59 and CD55 in WT cells were set as 1. The relative values were calculated and are presented as the means ± SD from thre e independent experiments. *P*-values (one-tailed, student’s t-test) are shown. (F) Misfolded CD59 (EGFP-FLAG-CD59 (C94S)) interacted with calnexin in PDIA3-KO cells. Constructs expressing misfolded CD59 (EGFP-FLAG-CD59 (C94S)) or soluble misfolded CD59 (EGFP-FLAG-CD59 (C94S, G103*)) were transiently transfected into WT and PDIA3-KO cells. Three days after transfection, cells were collected and lysed with buffer containing 0.5% CHAPS. The lysates were immunoprecipitated with anti-FLAG affinity gel, and after washing, the precipitated proteins were released by the addition of SDS sample buffer. The input (2.5% of total protein) and immunoprecipitated fractions (IP) were analyzed by immunoblotting with the indicated antibodies.

Nuclear magnetic resonance analysis and in vitro experimental results have indicated that the P-domain of calnexin is important for its association with the protein disulfide isomerases ERp57 (Frickel et al., 2002; M. R. Leach, Cohen-Doyle, Thomas, & Williams, 2002; Pollock et al., 2004) and ERp29 (Kozlov et al., 2017) through the tip of its arm domain. Furthermore, tryptophan at position 342 (W342), aspartic acid residues at positions 343 (D343) and 347 (D347), and glutamic acid at position 351 (E351) of calnexin are thought to be important for this association (Kozlov et al., 2017; Pollock et al., 2004). To assess the importance of these residues in the association of calnexin with ERp57 and ERp59, single (D343A, E351A) or double mutant calnexin constructs (W342A D343A, D343A E351A, D347A E351A) were constructed to disrupt the interaction. In cells transfected with these mutants, the PI-PLC sensitivity of CD59 was restored, whereas some of the constructs affected in their protein stability (Figure 3B and C), suggesting that calnexin participates in GPI-inositol deacylation independent of its binding with ERp57. To assess this possibility, *PDIA3*, which encodes ERp57, was knocked out in HEK293 cells, and the PI-PLC sensitivity of GPI-APs was analyzed. Interestingly, the PI-PLC sensitivity of both CD59 and CD55 was increased in PDIA3-KO cells compared to that observed in the parental WT cells (Figure 3D and E), suggesting that GPI-inositol deacylation occurred more efficiently in the absence of ERp57.

Calnexin was observed to associate with CD59 in an N-glycan- and GPI-dependent manner (Figure 3-figure supplement C), as previously observed (Liu et al., 2018). ERp57 weakly co-precipitated with CD59 and misfolded CD59, whereas interactions with other ER chaperones, such as calreticulin and BiP, were not detected with any CD59 construct (Figure 3 supplement C), indicating that the N-glycan and membrane-associated domain of GPI-APs are important for calnexin interaction. To assess whether ERp57 is required for calnexin to bind CD59, we expressed misfolded CD59 (C94S) and misfolded and soluble CD59 (C94S G103stop) in WT and PDIA3-KO cells and analyzed their ability to bind calnexin. Calnexin was co-precipitated with misfolded CD59 (C94S) but not misfolded and soluble CD59 (C94S G103stop). Furthermore, knockout of *PDIA3* did not affect the interaction between misfolded CD59 (C94S) and calnexin (Figure 3F), suggesting that calnexin binds with misfolded GPI-APs in an ERp57-independent manner.

### Disruption of the calnexin/calreticulin cycle leads to the exposure of misfolded GPI-APs on the plasma membrane

Under steady state conditions, the majority of misfolded prion proteins localize in the ER (Satpute-Krishnan et al., 2014). Because it appears that calnexin retains misfolded GPI-APs in the ER until their correct folding, we investigated the role of calnexin in the ER retention of GPI-APs by analyzing the localization of misfolded GPI-APs in the absence of the calnexin/calreticulin cycle. In HEK293 cells, EGFP-FLAG-tagged CD59 (C94S) was primarily localized in the ER (Figure 4A, top). In CANX-KO and CALR-KO cells, CD59 (C94S) was retained in the ER, similar to WT cells. In contrast, in CANX&CALR-DKO cells, CD59 (C94S) was not retained in the ER but rather localized at the plasma membrane (Figure 4A, bottom). Quantification of CD59 (C94S) co-localization with the ER marker was performed using multiple images (N=28 for WT cells; N=25 for CANX&CALR-KO cells) (Figure 4D). The Pearson’s correlation coefficient for EGFP-FLAG-CD59 (C94S) versus mRFP-KDEL changed from 0.715±0.05 (mean±SD) in WT cells to 0.176±0.04 in CANX&CALR-KO cells. Furthermore, another misfolded GPI-AP, EGFP-FLAG-CD55 (C81A), was also expressed on the surface of CANX&CALR-DKO cells (Figure 4-figure supplement A).

**Figure 4.**
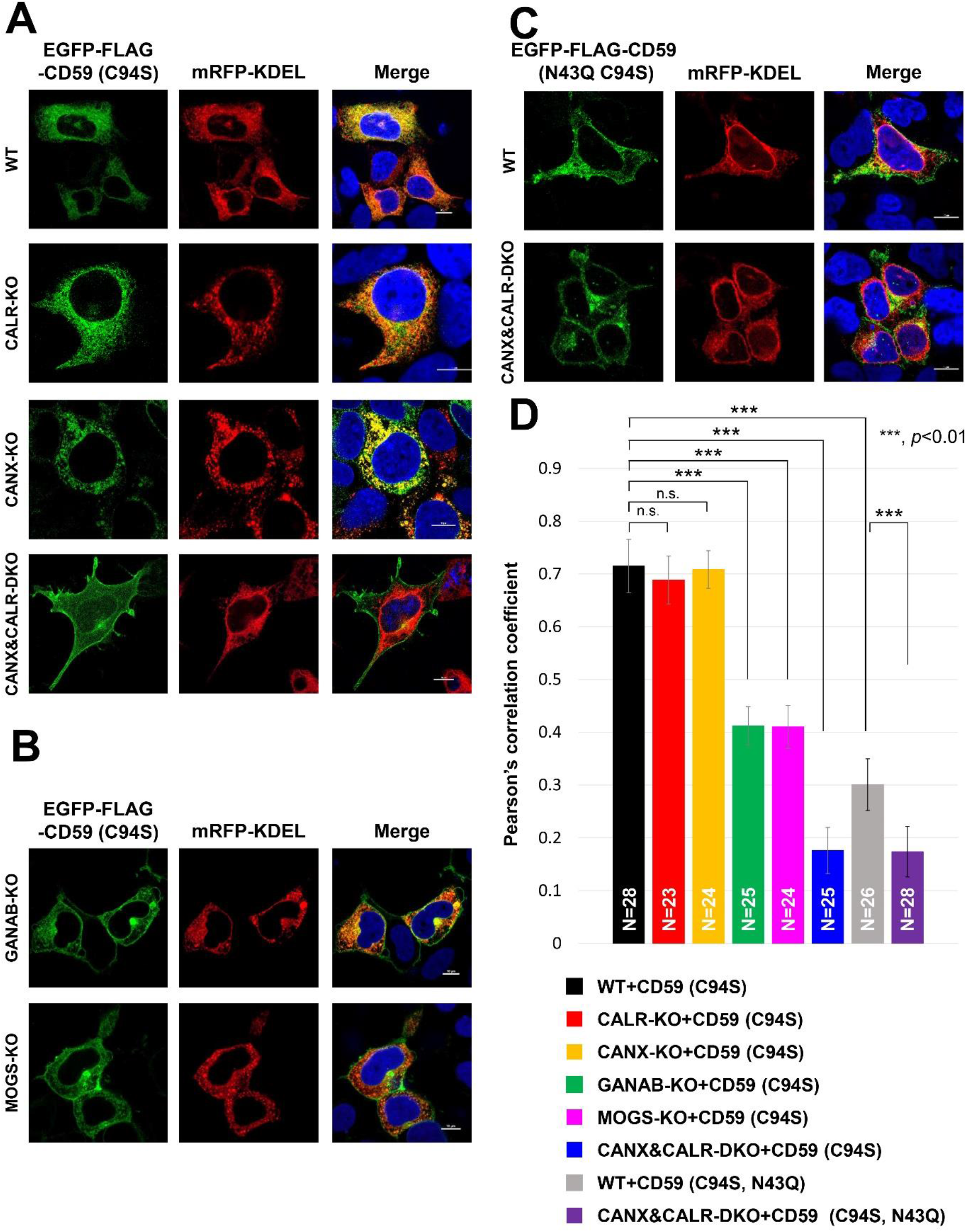
N-glycan-dependent calnexin/calreticulin cycle mediates the efficient ER retention of misfolded GPI-APs. (A-C) Localization of misfolded CD59 (EGFP-FLAG-CD59 (C94S)) was analyzed in WT, CANX-KO, CALR-KO, and CANX&CALR-DKO cells (A) and in MOGS-KO and GANAB-KO cells (B). Localization of misfolded and non-N-glycosylated CD59 (EGFP-FLAG-CD59 (C94S, N43Q)) was performed as described in (C). mRFP-KDEL was used as an ER marker. Three days after transfection, images were obtained using confocal microscopy. DAPI staining is shown as blue in merged images. Scale bar, 10 μm. (D) Pearson’s correlation coefficient values between EGFP-FLAG-CD59 (C94S) or (C94S, N43Q) and mRFP-KDEL were calculated using the ImageJ plugin JACoP. The data are presented as the means ± SD of the measurements. “N=” represents cell number used for the calculation. *P*-values (two-tailed, student’s t-test) are shown.

Since calnexin/calreticulin recognizes monoglucosylated N-glycan structures, we next analyzed the localization of misfolded CD59 in cells defective in glucose trimming. In MOGS-KO or GANAB-KO cells, the majority of N-glycan structures on proteins in the ER are retained as Glc_3_Man_9_GlcNAc_2_ and Glc_2_Man_9_GlcNAc_2_, which are not recognized by calnexin. In MOGS-KO and GANAB-KO cells, fractions of CD59 (C94S) were expressed on the cell surface (Figure 4B), supporting that misfolded GPI-APs lose their ability to interact with calnexin, causing their expression on the cell surface. The Pearson’s correlation coefficient analysis showed localization of EGFP-FLAG-CD59 (C94S) was changed from the ER in MOGS-KO and GANAB-KO cells. These results indicate that the calnexin/calreticulin cycle mediates the ER retention of misfolded GPI-APs.

To verify whether N-glycan structures on GPI-APs are required for their ER retention, the localization of non-N-glycosylated misfolded CD59 (C94S, N43Q) and CD55 (C81A, N95Q) was assessed. The results showed that these proteins were not retained in the ER, but fractions of them were expressed on the cell surface, even in the WT cells (Figure 4C, D and Figure 4-figure supplement B). In CANX&CALR-DKO cells, non-N-glycosylated misfolded CD59 and CD55 were further expressed on the plasma membrane (Figure 4C and Figure 4-figure supplement B). These results are consistent with the co-IP results showing that non-N-glycosylated misfolded CD59 lost its ability to interact with calnexin such that it could not be retained in the ER.

The above results suggest that calnexin retains misfolded GPI-APs in the ER through its binding to N-glycans on these proteins. To further investigate this possibility, we assessed the localization of misfolded CD59 in CANX&CALR-DKO cells expressing lectin-deficient calnexin. The lectin-deficient calnexin could not rescue the phenotype of CANX&CALR-DKO cells (Figure 4-figure supplement C), whereas mutant calnexin defective in ERp57 interaction could rescue and retain misfolded CD59 in the ER (Figure 4-figure supplement D). These results provide evidence that the lectin domain of calnexin is responsible for the ER retention of N-glycosylated GPI-APs.

To assess the specificity of the localization changes in CANX&CALR-DKO cells, a transmembrane form of CD59 (EGFP-FLAG-CD59-TM (C94S)), in which the GPI-attachment signal was replaced with a transmembrane domain of CD46, was constructed, and its localization was analyzed in WT and CANX&CALR-DKO cells. The transmembrane form of misfolded CD59 was localized in the ER in WT cells, whereas the proteins were still retained in the ER of CANX&CALR-DKO cells (Figure 4-figure supplement E), suggesting that the GPI-anchor is responsible for the change in misfolded protein localization in CANX&CALR-DKO cells.

### Misfolded and inositol-acylated GPI-APs are exposed on the plasma membrane in CANX&CALR-DKO cells

The results of a previous study confirmed that misfolded prions resulting from acute ER stress are released to the secretory pathway and can reach the plasma membrane (Satpute-Krishnan et al., 2014). In the present study, by treating cells with 100 nM thapsigargin, the localization of misfolded CD59 (C94S) began to shift from the ER to the cell surface (Figure 5A and B). After thapsigargin (TG) treatment, misfolded CD59 was expressed on the cell surface, whereas calnexin was intracellularly retained (Figure 5-figure supplement A).

**Figure 5.**
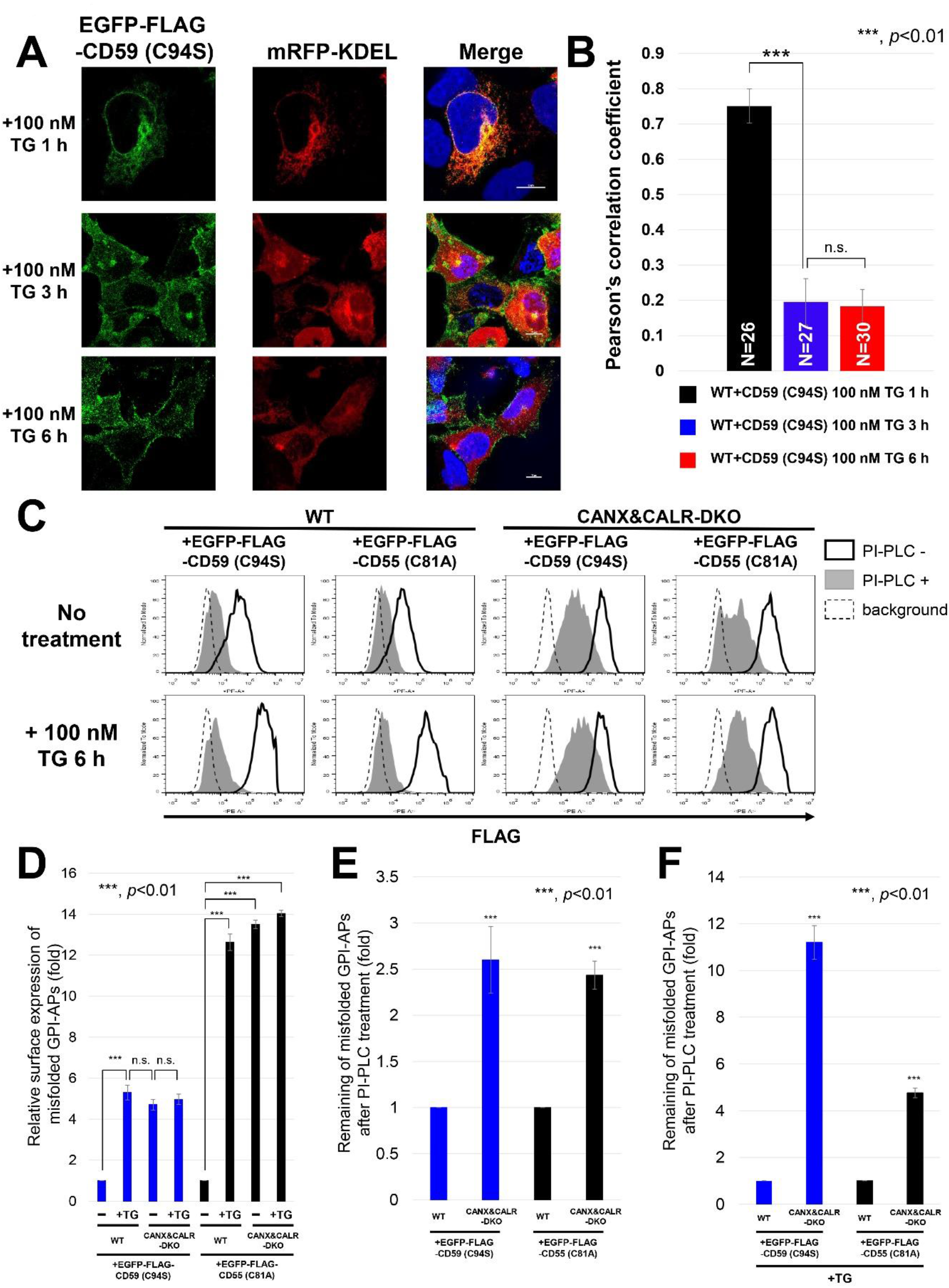
Misfolded and inositol-deacylated GPI-APs are exposed on the plasma membrane under acute ER stress conditions. (A and B) EGFP-FLAG-CD59 (C94S) and mRFP-KDEL constructs were transiently transfected into WT cells. Three days after transfection, the cells were treated with 100 nM thapsigargin (TG) for 1, 3 and 6 h. The cells were fixed and imaged by confocal microscopy (A). Scale bar, 10 μm. Pearson’s correlation coefficient values between EGFP-FLAG-CD59 (C94S) and mRFP-KDEL were calculated using the ImageJ plugin JACoP (B). The data are presented as the means ± SD of the measurements. “N=” represents cell number used for the calculation. (C) EGFP-FLAG-CD59 (C94S) or EGFP-FLAG-CD55 (C81A) constructs together with a BFP-expressing plasmid were transiently transfected into WT and CANX&CALR-DKO cells. Three days after transfection, the cells were incubated with or without 100 nM TG for 6 h and then treated with or without PI-PLC. The surface expression of EGFP-FLAG-CD59 (C94S) and EGFP-FLAG-CD55 (C81A) was analyzed by flow cytometry. Cells showing the same BFP intensities were gated for the same transfection level. (D) Surface expression of misfolded CD59 (EGFP-FLAG-CD59 (C94S)) or CD55 (EGFP-FLAG-CD55 (C81A)) in WT or CANX&CALR-DKO cells with or without TG treatment for 6 h shown in Figure 5C (heavy solid line) were plotted. The data are presented as the means ± SD of three independent measurements. *P*-values (one-tailed, student’s t-test) are shown. (E and F) After PI-PLC treatment, surface misfolded CD59 (EGFP-FLAG-CD59 (C94S)) and CD55 (EGFP-FLAG-CD55 (C81A)) in WT or CANX&CALR-DKO cells without (E) or with (F) TG treatment for 6 h shown in Figure 5C (gray shaded) were plotted. The data are presented as the means ± SD of three independent measurements. *P*-values (one-tailed, student’s t-test) are shown.

To confirm the specificity of this phenomena that were observed in misfolded GPI-APs, we analyzed the localization of an ERAD substrate, HLA1-1-147 (Zhang, Xu, Liu, & Ye, 2015), and that of the transmembrane form of misfolded CD59 (C94S)-TM with or without ER stress induction. Both HLA1-1-147 and CD59 (C94S)-TM remained localized in the ER, even after cells were treated with thapsigargin for 6 h (Figure 5-figure supplement B and C), suggesting that the observed localization changes are specific for GPI-APs.

By induction of acute ER stress, misfolded GPI-APs are released into the secretory pathway and can be expressed on the plasma membrane. Flow cytometry results showed that fractions of misfolded CD59 and CD55 were expressed on the cell surface in WT cells without the ER stress, while this surface expression was significantly increased after treatment with thapsigargin for 6 h (by 5.3- and 12.6-fold for CD59 and CD55, respectively) (Figure 5C, left panels, heavy solid line and Figure 5D). The misfolded CD59 and CD55 on the surface of WT cells showed PI-PLC sensitivity (Figure 5C, left panels, gray peak). On the other hand, on the surface of CANX&CALR-DKO cells, the levels of misfolded CD59 and CD55 were 4.7- and 13.5-fold higher than those observed in WT cells without thapsigargin treatment, respectively (Figure 5C, right panels, heavy solid line and Figure 5D). By PI-PLC treatment, misfolded CD59 and CD55 showed PI-PLC resistance (2.6 and 2.4 times higher) in CANX&CALR-DKO cells, compared with WT cells (Figure 5C, top panels, gray peak and Figure 5E). ER stress induction by thapsigargin further increased the PI-PLC resistance of CD59 and CD55 in CANX&CALR-DKO cells compared with that observed in WT cells (11.2- and 4.7-fold higher, respectively) (Figure 5C, bottom panels, gray peak and Figure 5F). These results indicate that misfolded GPI-APs in the ER of WT cells are already inositol-deacylated, and express on the cell surface by ER stress induction. On the other hand, misfolded GPI-APs in CANX&CALR-DKO cells are expressed regardless of acute ER stress, parts of whose GPI-moieties are not processed. The difference in PI-PLC sensitivity of GPI-APs between CANX&CALR-DKO cells and thapsigargin-treated WT cells suggests that ER retention time is important for efficient GPI-inositol deacylation. In the WT cells, misfolded GPI-APs were retained in the ER for a long time, providing them sufficient time to interact with the GPI-inositol deacylase PGAP1. In contrast, in CANX&CALR-DKO cells, misfolded GPI-APs could not be retained in the ER.

### Alkaline phosphatase activity is greatly decreased in CANX&CALR-DKO cells compared to that observed in WT cells

Given the results showing that misfolded GPI-APs were expressed on the plasma membrane in CANX&CALR-DKO cells, we hypothesized that misfolded/unfolded endogenous GPI-APs are expressed on the plasma membrane in CANX&CALR-DKO cells under normal conditions. Alkaline phosphatases are GPI-APs, for which there are four types, including tissue non-specific, intestinal, placental, and placental-like typestypes in human. Among these different types, the expression of the placental alkaline phosphatase (ALPP, contain 2 N-glycosylation sites) is the highest in HEK293 cells (Figure 6-figure supplement). The surface expression of endogenous ALPP was almost the same in WT and CANX&CALR-DKO cells (Figure 6A, heavy solid lines and Figure 6B). The ALPP showed 2.3-fold higher PI-PLC resistance in CANX&CALR-DKO cells compared to that observed in WT cells (Figure 6A, gray peaks and Figure 6C). We then assessed the ALP activity on the cell surface and compared with WT cells, the activity in CANX&CALR-DKO cells was significantly decreased by 60% (Figure 6D). These results indicate that GPI-APs that are not fully functional are expressed on the plasma membrane in CANX&CALR-DKO cells.

**Figure 6.**
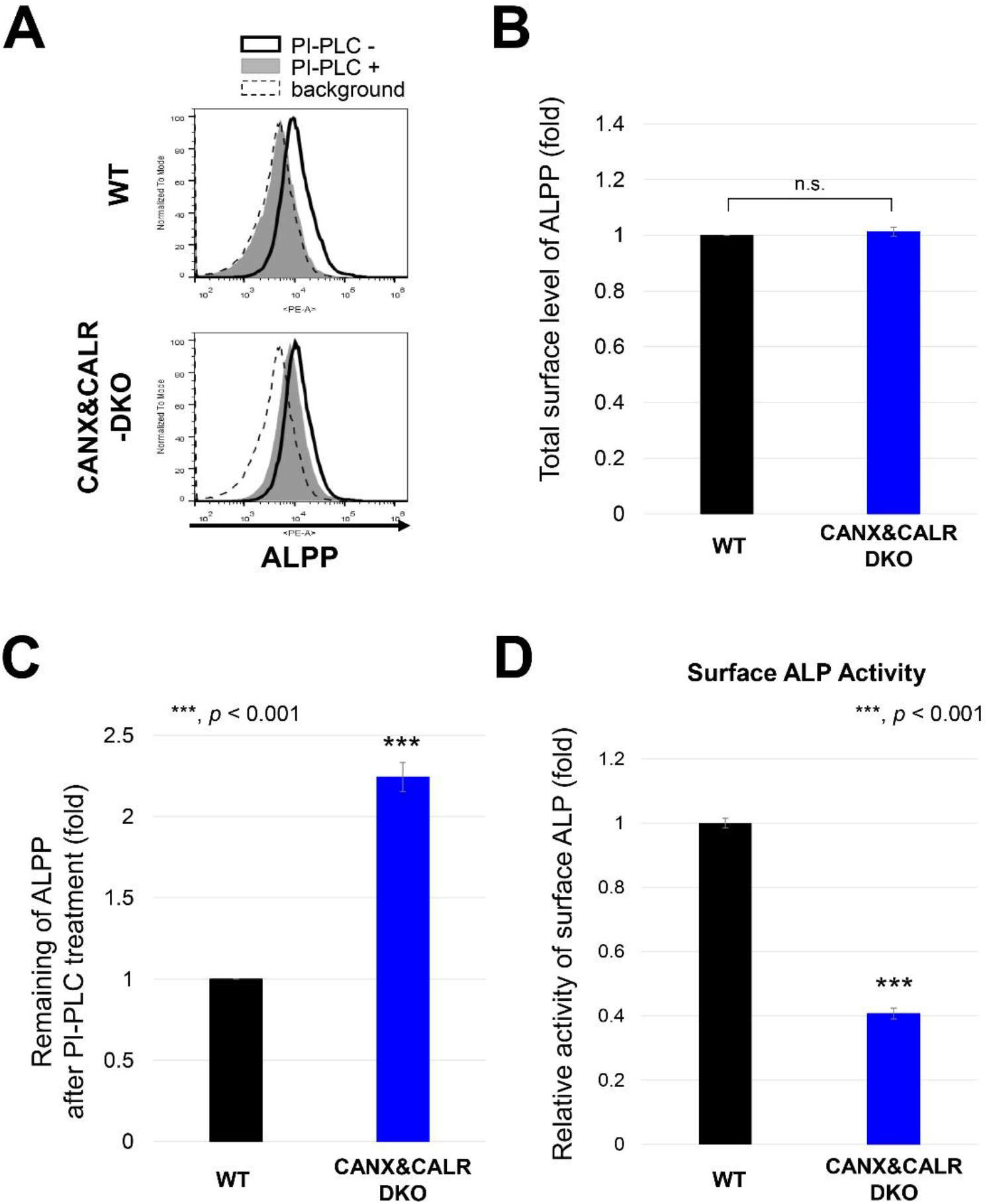
ALPP activity decreases in CANX&CALR-DKO cells. (A – C) Sensitivity of PI-PLC to placental alkaline phosphatase (ALPP) in WT and CANX&CALR-DKO cells was analyzed by flow cytometry (A). An anti-ALPP antibody was used as the primary antibody. The dashed lines show the background of the isotype control. The surface expression of ALPP in WT or CANX&CALR-DKO cells was plotted (B). The data are presented as the means ± SD of three independent measurements. The levels of ALPP remaining after PI-PLC treatment are plotted (C). The remaining ALPP values in WT cells were set as 1. The relative values were calculated and are presented as the means ± SD from three independent experiments. P-values (one-tailed, student’s t-test) are shown. (D) The relative activity of surface ALP in WT and CANX&CALR-DKO cells are shown. The activity of surface ALP was measured as described in the Materials and Methods. The data are presented as the means ± SD of three independent measurements. P-values (two-tailed Student’s t-test) are shown.

### Sufficient ER retention time is required for efficient inositol deacylation by PGAP1

To confirm whether sufficient ER retention time is required for GPI-inositol deacylation, a doxycycline (DOX)-inducible VSVG^ts^-FLAG-GFP-GPI (VFG-GPI) reporter system (Maeda et al., 2008) was used. VFG-GPI is a temperature-sensitive glycoprotein, and the reporter causes misfolding at 40°C due to a mutation in the VSVG portion and is retained in the ER, whereas it is folded at 32°C and transported to the plasma membrane (Fujita et al., 2009; Fujita et al., 2011; Hirata et al., 2015). Since CANX&CALR-DKO cells cannot retain misfolded GPI-APs such as CD59 and CD55 in the ER under normal culture conditions (37°C), the localization of VFG-GPI in CANX&CALR-DKO cells at 40°C was assessed. VFG-GPI was localized in the ER in WT, CANX&CALR-DKO, and CANX-rescued CANX&CALR-KO cells (Figure 7A) under 40°C.

**Figure 7.**
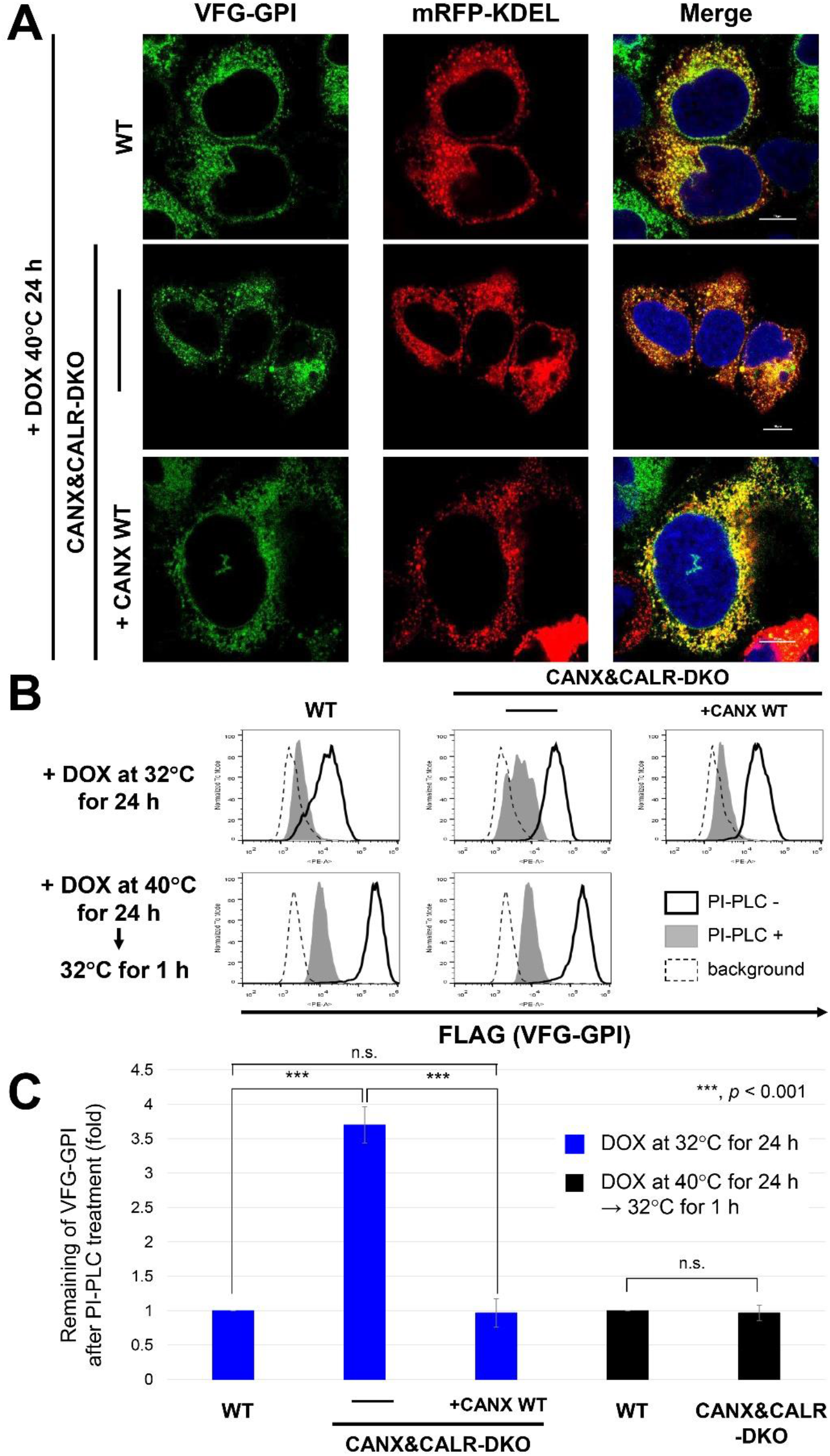
ER retention time regulates GPI-inositol deacylation. (A) Localization of VSVGts-FLAG-GFP-GPI (VFG-GPI) at 40°C in WT, CANX&CALR-DKO, and CANX&CALR-DKO+CANX WT cells. The construct expressing mRFP-KDEL, an ER marker, was transfected into WT, CANX&CALR-DKO and CANX&CALR-DKO+CANX WT cells. At 36 h after transfection, cells were incubated with 1 μg/ml doxycycline at 40°C for 24 h to induce VFG-GPI expression. Subsequently, the cells were quickly fixed with 4% paraformaldehyde and then imaged by confocal microscopy. Scale bar, 10 μm. (B) HEK293FF6WT, CANX&CALR-DKO and CANX&CALR-DKO+CANX WT cells were incubated with with 1 μg/ml doxycycline at 32°C for 24 h (upper panels). Alternatively, cells were incubated with 1 μg/ml doxycycline at 40°C for 24 h followed by a 32°C incubation for 1 h (lower panels). After induction, the cells were treated with or without PI-PLC, and surface VFG-GPI was stained with an anti-FLAG antibody and analyzed by flow cytometry. The region of same GFP intensity was gated to normalize VFG-GPI expression. (C) Remaining of VFG-GPI after PI-PLC treatment in (B) were plotted. The values in WT cells were set as 1, and relative values in CANX&CALR-DKO were plotted. The data are presented as the means ± SD of three independent measurements. *P*-values (one-tailed, student’s t-test) are shown.

We then analyzed surface-expressed VFG-GPI under different conditions. When VFG-GPI expression was induced at 32°C, VFG-GPI in WT and CANX-rescued CANX&CALR-DKO cells was sensitive to PI-PLC (Figure 7B, top), whereas partial VFG-GPI resistance to PI-PLC was detected in CANX&CALR-DKO cells (3.6-fold higher than that measured for WT cells) (Figure 7B, top panels, middle and Figure 7C, left), which is consistent with that observed for other GPI-APs.

We next analyzed VFG-GPI using a chase experiment. VFG-GPI was expressed at 40°C to accumulate in the ER, after which the temperature was shifted to 32°C to allow its transport to the plasma membrane. Under these condition, surface VFG-GPI in CANX&CALR-DKO became sensitive to PI-PLC cleavage, similar to that observed in WT cells (Figure 7B, bottom panels and Figure 7C, right). These results proved that sufficient ER retention time is necessary for GPI-inositol deacylation.

## Discussion

Both calnexin-mediated ER quality control and GPI-inositol deacylation by PGAP1 are performed in the ER. The calnexin/calreticulin cycle is essential for N-glycosylated protein folding (Tannous et al., 2015), and GPI-inositol deacylation is required for the efficient sorting of GPI-APs into transport vesicles and their subsequent transport from ER to the Golgi apparatus (Fujita & Kinoshita, 2012; Muñiz & Riezman, 2016). In the present study, we confirmed that calnexin not only contributes to protein folding but also GPI-inositol deacylation, playing dual roles in the maturation of GPI-APs in the ER.

In WT cells, calnexin was observed to retain GPI-APs in the ER until achieving correct folding by directly binding to the N-glycans on the GPI-APs (Figure 8, top). The results of our previous study showed that PGAP1 associates with calnexin (Liu et al., 2018). Interestingly, the temporal ER retention of GPI-APs and their association with PGAP1 mediated by calnexin promoted the efficient cleavage of acyl chains from GPI-anchors. Correctly folded and inositol deacylated GPI-APs are further processed by PGAP5 and recognized by p24 cargo receptors to become incorporated into transport vesicles. In CANX&CALR-DKO cells, newly synthesized GPI-APs could not be retained in the ER, and fractions of GPI-APs were transported without correct folding and GPI processing (Figure 8, middle). Since structural remodeling of GPI-APs is crucial for their binding to p24 cargo receptors, inositol-acylated GPI-APs would exit from the ER through bulk flow pathway. Since ER quality control systems are impaired in CANX&CALR-DKO cells, both correctly folded and misfolded GPI-APs are expressed on the cell surface. The observation that more than 90% of GPI-APs are N-glycosylated (Liu et al., 2018) indicates that calnexin-dependent GPI-anchor maturation is a commonly used pathway. In addition, calnexin-independent GPI-inositol deacylation occurs for non-N-glycosylated GPI-APs. Other molecular chaperones such as BiP and protein disulfide isomerases, which possess ER-retention signals, may be involved in the folding and GPI-processing of non-N-glycosylated proteins. When misfolded GPI-APs are expressed under basal conditions, they are retained in the ER by calnexin. During the long ER retention time, an acyl-chain on the GPI-inositol is cleaved by PGAP1 (Figure 8, bottom). Since the GPI moiety of misfolded ER-resident GPI-APs signals that they are ready to exit, they would be quickly transported from the ER via acute ER stress induction.

**Figure 8.**
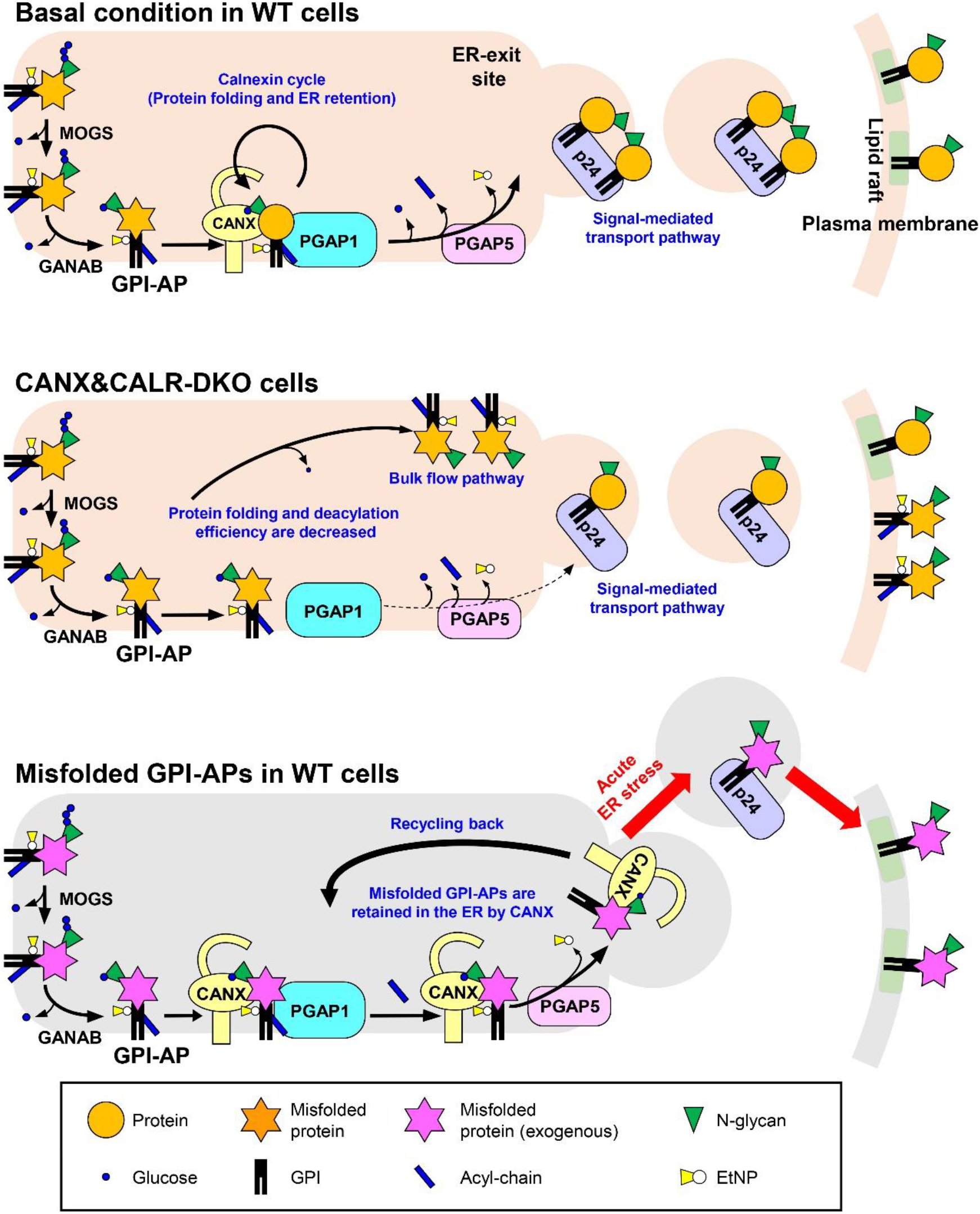
Summary models for folding and inositol deacylation of GPI-APs regulated by calnexin/calreticulin. Top: Under normal conditions, N-glycans and GPI are transferred to the newly synthesized GPI-APs. After glucose trimming by α-glucosidase I and II, the N-glycan structure becomes Glc_1_Man_9_GlcNAc_2_, which is specifically recognized by calnexin/calreticulin. By directly binding with calnexin, immature GPI-APs enter the ER quality control system and finally become mature GPI-APs. In addition, calnexin retains GPI-APs in the ER and associates with PGAP1, which is required for the efficient GPI-inositol deacylation. After the GPI moiety is remodeled by PGAP1 and PGAP5, and GPI-APs are transported from the ER to the Golgi by the cargo receptors, p24 family of proteins. Middle: In CANX&CALR-DKO cells, there is no calnexin/calreticulin-dependent ER quality control system and ER retention system. GPI-APs exit the ER with incomplete protein folding and GPI remodeling. Bottom: In WT cells, under basal conditions, misfolded GPI-APs are retained in the ER by calnexin. During the long time being retained in the ER, an acyl-chain on the GPI-inositol is removed by PGAP1. Once acute ER stress is induced, deacylated and misfolded GPI-APs are quickly transported by p24 family members and exposed on the plasma membrane.

Calnexin/calreticulin interact with newly synthesized proteins in an N-glycan-dependent manner. However, once high molecular weight aggregates of the misfolded protein are generated, calnexin can interact with them in an N-glycan-independent manner (Cannon, Hebert, & Helenius, 1996; Thammavongsa, Mancino, & Raghavan, 2005). N-glycan-dependent calnexin binding is a mechanism to protect against misfolded GPI-APs becoming insoluble aggregates, while N-glycan-independent binding will lead to the degradation of misfolded proteins (Seckler & Jaenicke, 1992; Swanton, High, & Woodman, 2003). Calnexin binding to GPI-APs was observed to occur in an N-glycan-dependent and PDI-independent manner. In PDIA3-KO cells, GPI-APs showed greater PI-PLC sensitivity, suggesting that ERp57 is dispensable for their binding and that calnexin could retain GPI-APs for longer time in the ER to allow their correct folding and GPI-processing.

Rapid ER stress-induced export (RESET) is a degradation pathway for misfolded GPI-APs under acute ER stress conditions (Satpute-Krishnan et al., 2014). Misfolded GPI-APs are retained in the ER under normal conditions. However, through the induction of acute ER stress, misfolded GPI-APs are transported from the ER to the plasma membrane and degraded in lysosomes. Our results are consistent with this process. Misfolded GPI-APs are recognized by calnexin, which is dependent on N-glycan for retention in the ER. We demonstrated that interaction of GPI-APs with calnexin contributes to GPI-anchor processing by PGAP1. Recently, misfolded prions have been reported to be expressed on the plasma membrane in association with calnexin and p24 family proteins by the induction of acute ER stress (Zavodszky & Hegde, 2019). In the present study, we showed that misfolded GPI-APs were stably expressed on the cell surface in the absence of calnexin and calreticulin, indicating that at least calnexin is not required for the exit of GPI-APs from the ER under stress conditions. Disruption of the calnexin/calreticulin cycle leads to the exit of misfolded GPI-APs from the ER due to a lack of ER retention mechanisms. In the RESET pathway, misfolded GPI-APs are subsequently endocytosed and delivered to lysosomes. In CANX&CALR-DKO cells, misfolded GPI-APs were stably expressed on the cell surface. In future studies, it would be worth identifying determinants for the endocytosis of misfolded GPI-APs.

Misfolded CD59 (C94S) and CD55 (C81A) have been shown to be the substrates for the RESET pathway (Satpute-Krishnan et al., 2014). When we replaced the GPI-attachment signal at the C-terminus to the transmembrane domain of CD46 (CD59 (C94S) TM) or used an ERAD substrate HLA1-1-147, their localization from the ER to the plasma membrane was not altered in CANX&CALR-DKO cells or under thapsigargin treatment. Therefore, it appears that the change in the localization of misfolded proteins in CANX&CALR-DKO cells is specific to GPI-APs. Since GPI-APs are tethered to the membrane through a phospholipid, their behaviors would be close to soluble proteins or phospholipids rather than transmembrane proteins. Some soluble ER-resident proteins having ER-retention signals have been reported to be secreted by the depletion of ER calcium (Trychta, Back, Henderson, & Harvey, 2018), which is a trigger of the unfolded stress response. It remains unclear how such ER-resident proteins are secreted in response to ER stress conditions. Thus, analysis of misfolded GPI-APs may allow for a better understanding of the associated mechanisms, which should be addressed in future studies.

## Supporting information

Key resources table

## Acknowledgements

We thank Drs Hideki Nakanishi, Ganglong Yang, Ning Wang (Jiangnan University) for their critical reading of the manuscript and comments. This work was supported by grants-in-aid from the National Natural Science Foundation of China (31770853), the Program of Introducing Talents of Discipline to Universities (111-2-06), National first-class discipline program of Light Industry Technology and Engineering (LITE2018-015), Top-notch Academic Programs Project of Jiangsu Higher Education Institutions, the International Joint Research Laboratory for Investigation of Glycoprotein Biosynthesis at Jiangnan University, and a grant for Joint Research Project of the Research Institute for Microbial Diseases, Osaka University.

## Supplemental Figures

**Figure 1-supplement.**
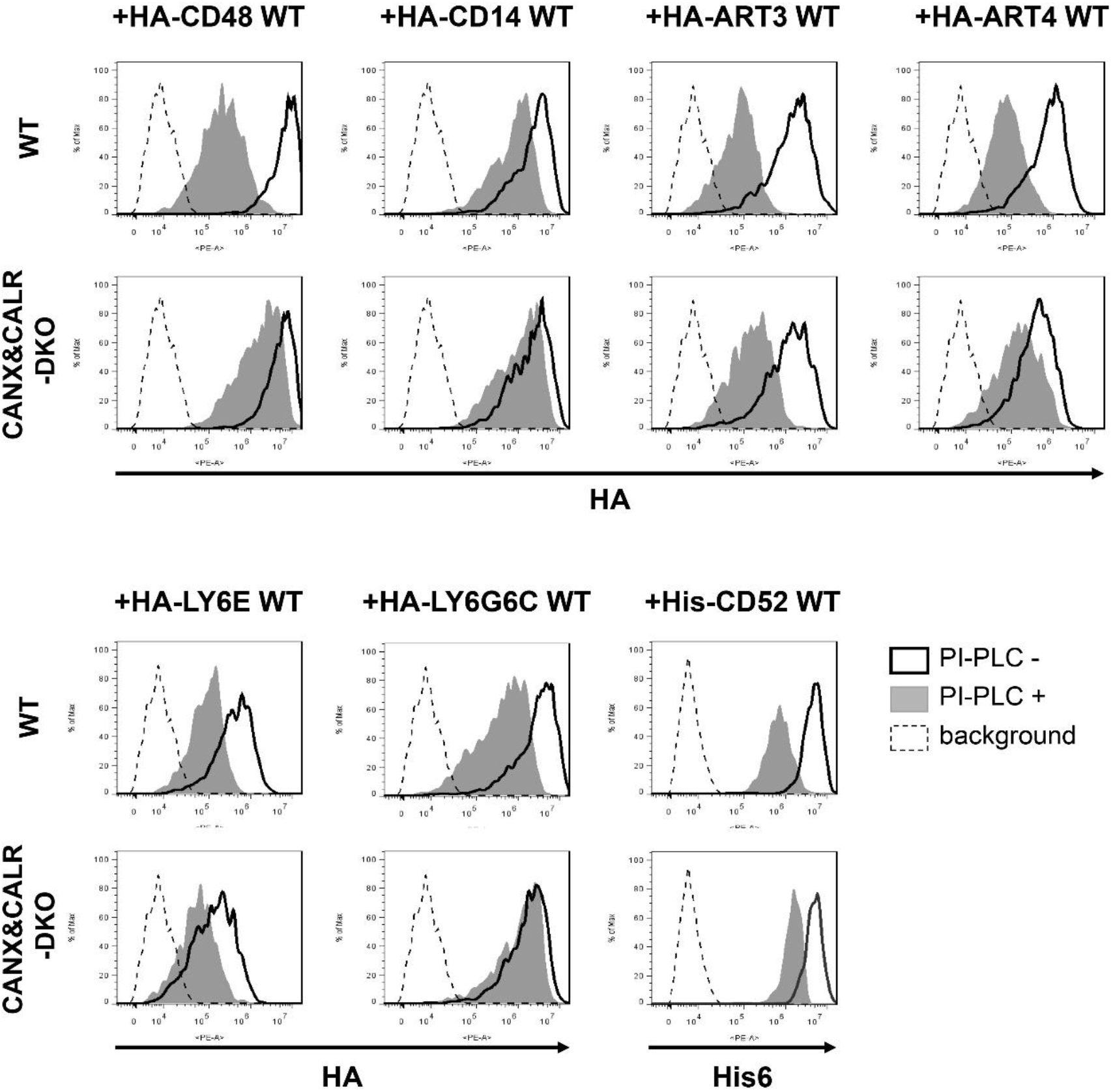
Calnexin/calreticulin-dependent GPI-inositol deacylation is commonly observed in N-glycosylated GPI-APs. HA-tagged or His6-tagged exogenous GPI-APs were expressed in WT and CANX&CALR-DKO cells, and their PI-PLC sensitivity was analyzed. The pME-HA-GPI-AP plasmid was transfected together with pME-BFP into WT and CANX&CALR-DKO cells. The plasmid pLIB2-His6-GPI-AP-IRES-BFP was transfected into HEK293T packaging cells, and retroviral vectors were infected into WT and CANX&CALR-DKO cells. Three days after transfection, cells were treated with or without PI-PLC. Anti-HA or anti-His6 were used as the primary antibodies for staining the surface proteins, which were analyzed by flow cytometry. The cells showing the same BFP intensities were gated. The values are shown in Figure 1C.

**Figure 3-supplement.**
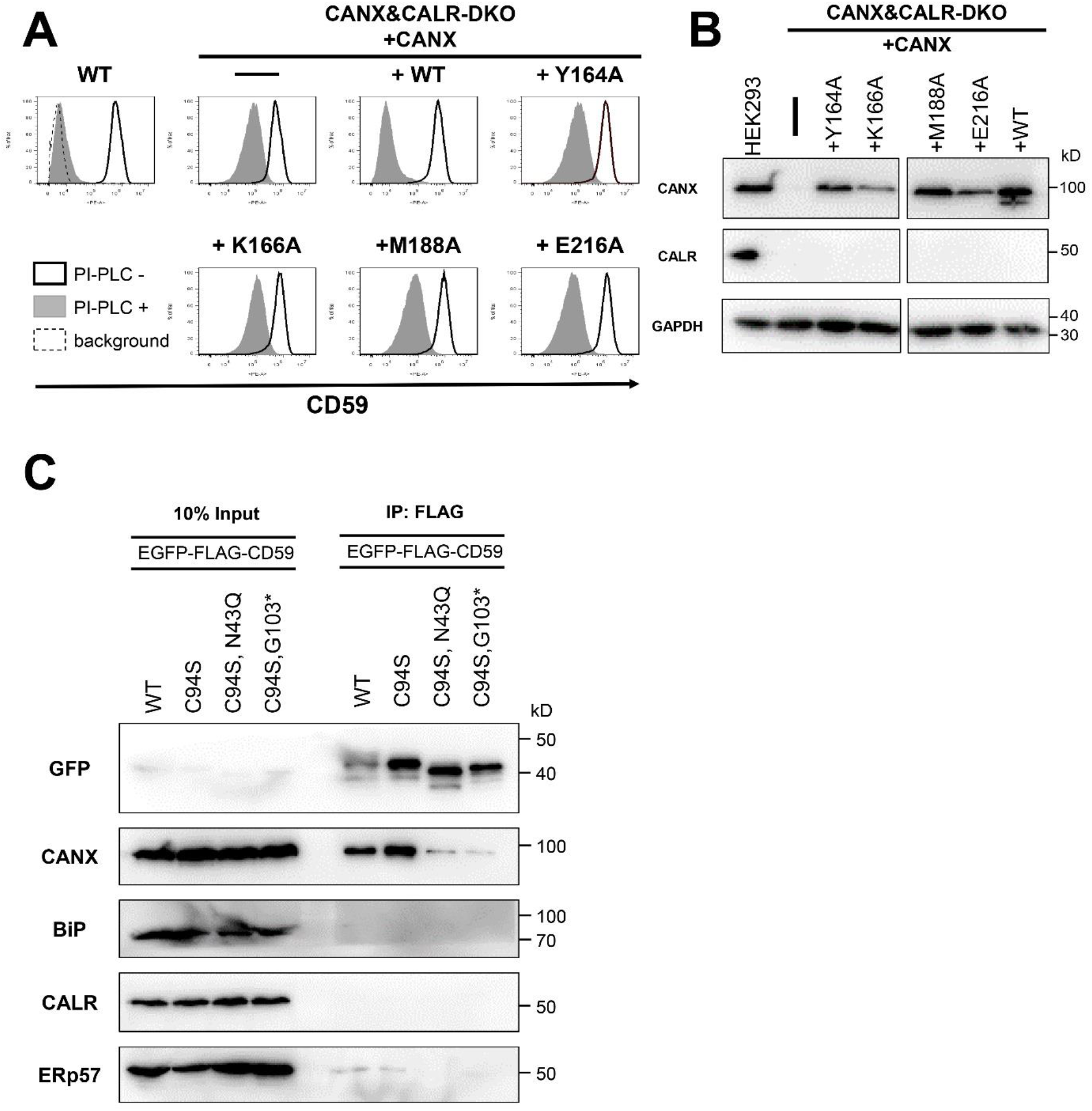
N-glycan-binding activity of calnexin is necessary for efficient GPI-inositol deacylation. (A and B) WT and CANX&CALR-DKO cells stably expressing WT CANX or CANX mutants defective in lectin activity (Y164A, K166A, M188A or E216A) were treated with or without PI-PLC. Surface CD59 was detected by flow cytometry (A) as described in Figure 1B. CANX and CALR from cell lysates in (A) were analyzed by immunoblotting (B). GAPDH was used as a loading control. (C) Cells were transiently transfected with constructs expressing WT EGFP-FLAG-CD59, misfolded CD59 (EGFP-FLAG-CD59 (C94S)), misfolded CD59 lacking GPI (EGFP-FLAG-CD59 (C94S, G103*)), or misfolded CD59 lacking an N-glycan (EGFP-FLAG-CD59 (C94S, N43Q)) and then were lysed with lysis buffer containing 0.5% CHAPS. The lysates were then subjected to immunoprecipitation (IP) with anti-FLAG affinity gel, and after washing, the precipitated proteins were released by the addition of SDS sample buffer. The input (10% of total protein) and immunoprecipitated fractions were analyzed by immunoblotting with the indicated antibodies.

**Figure 4-supplement.**
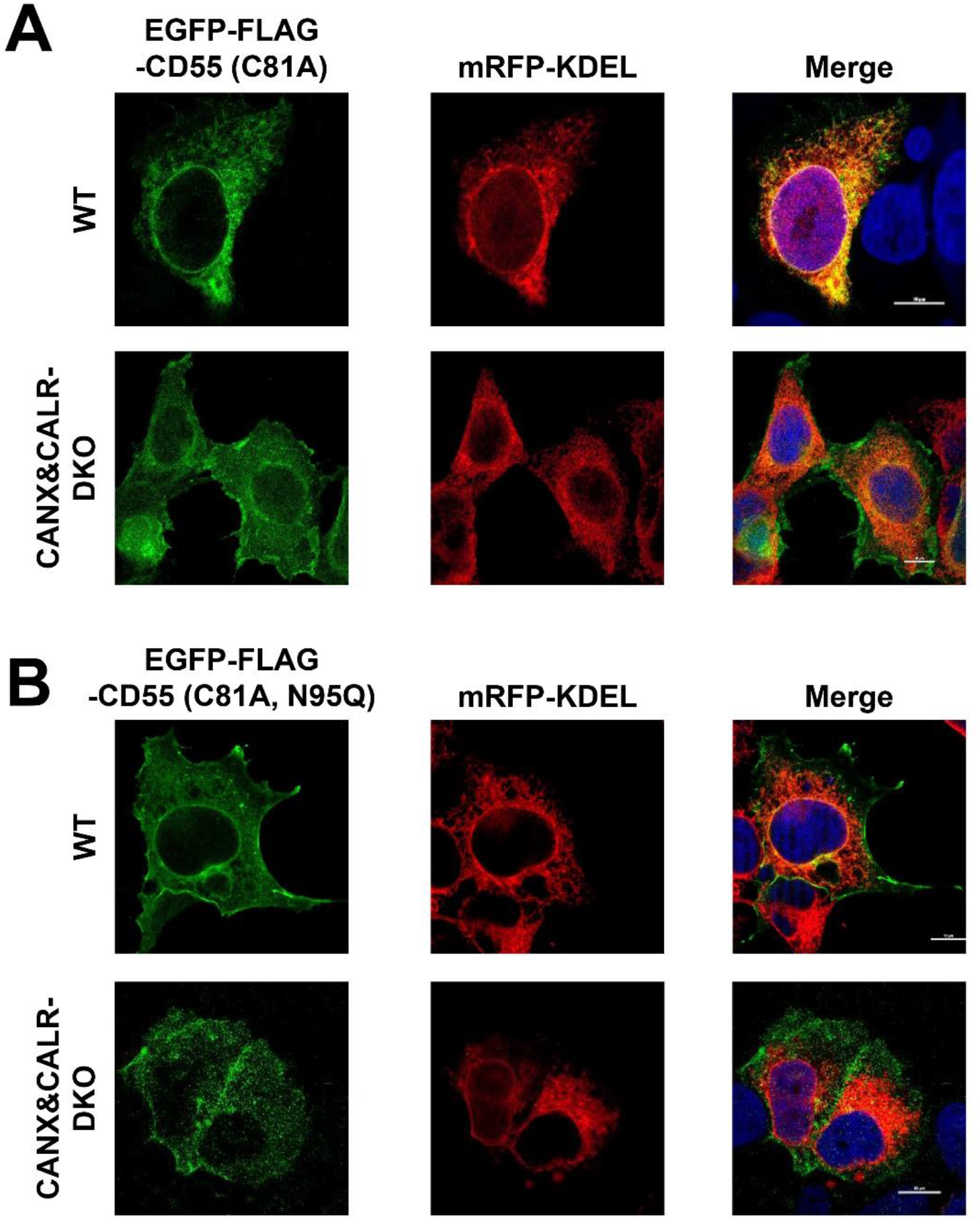
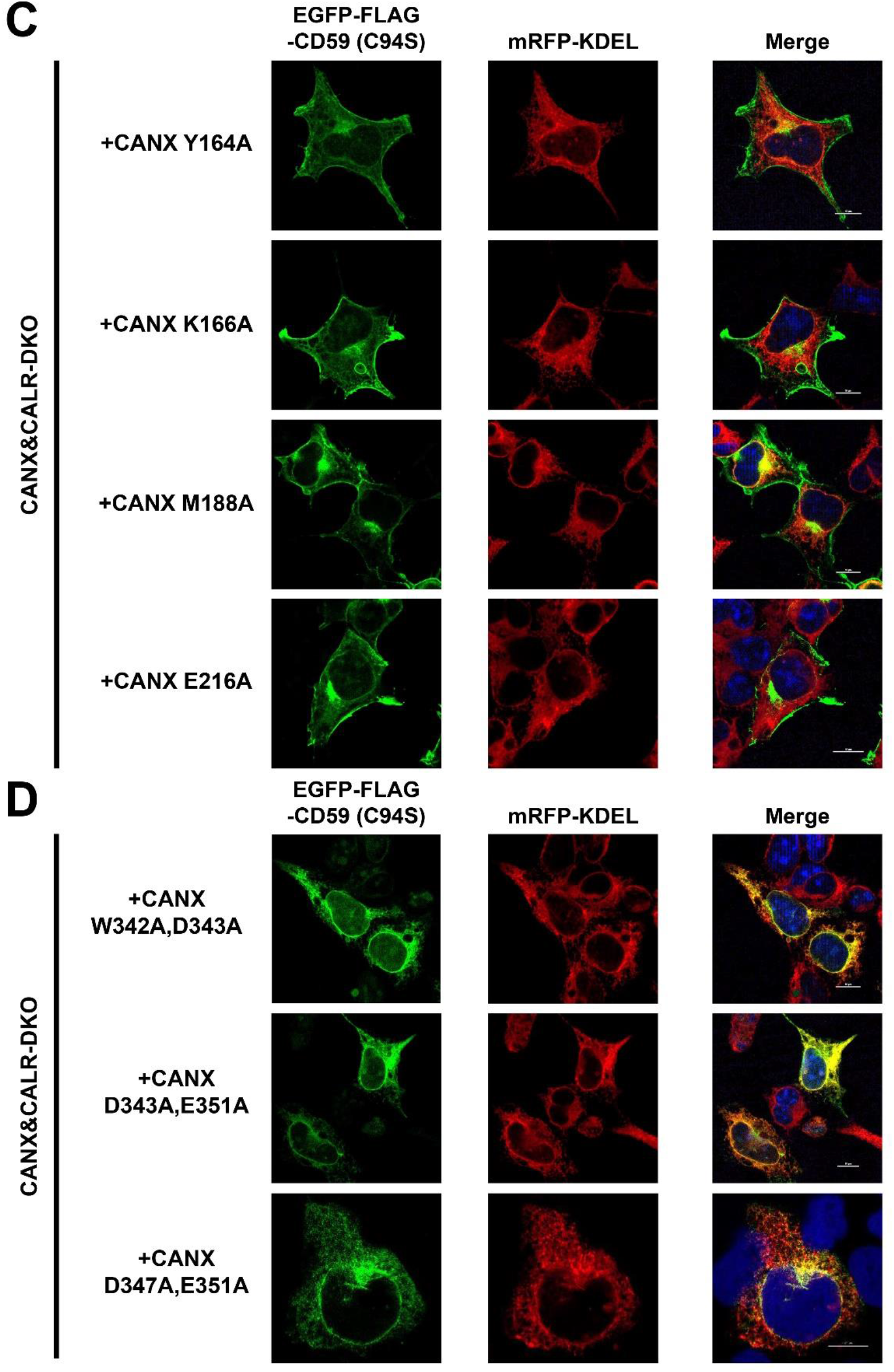
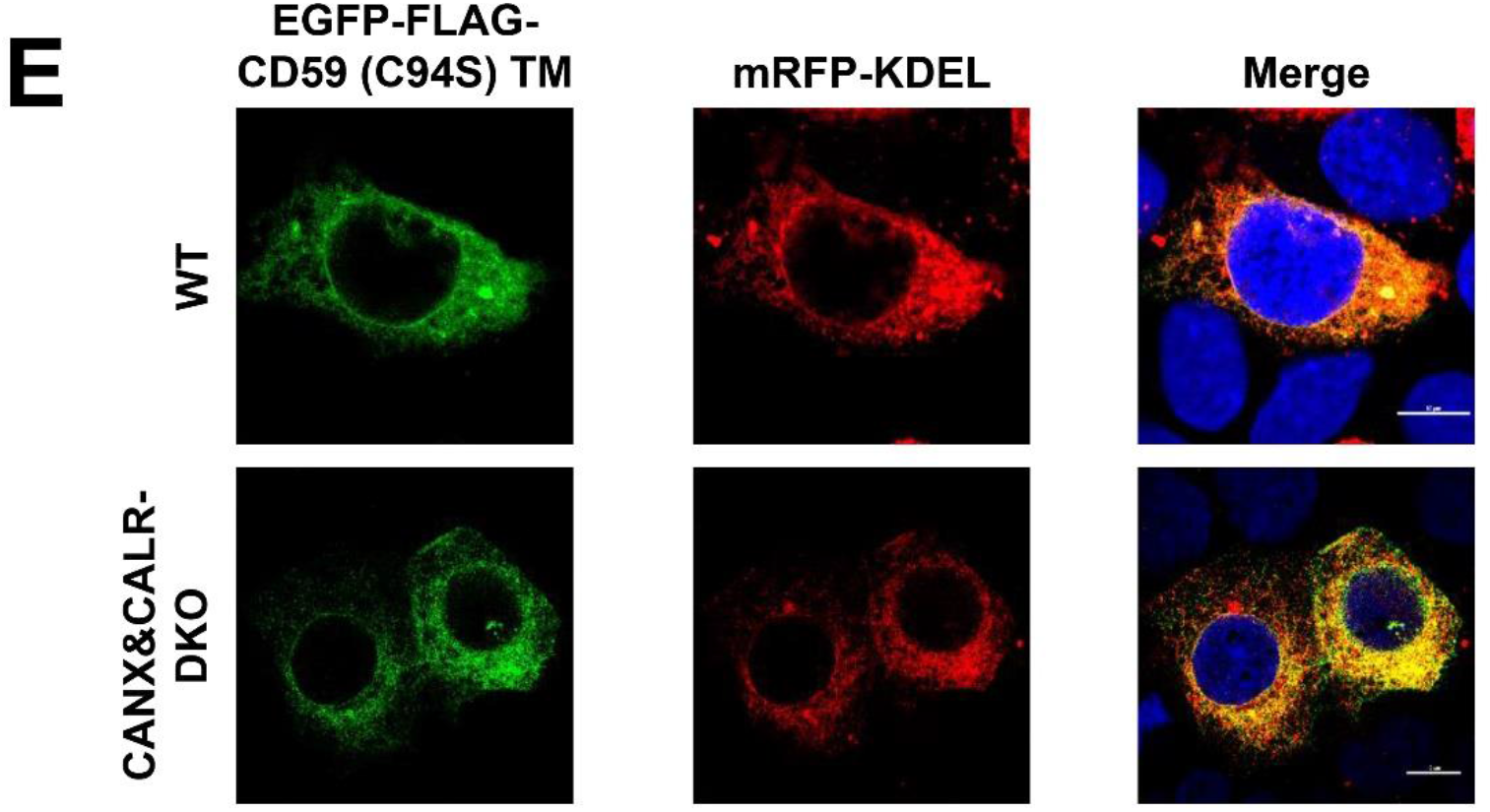
Localization of misfolded GPI-APs in CANX&CALR-DKO cells. (A and B) EGFP-FLAG-CD55 (C81A) or EGFP-FLAG-CD55 (C81A, N95Q) constructs were transiently transfected together with mRFP-KDEL into WT and CANX&CALR-DKO cells. The images were obtained using confocal microscopy at 3 days after transfection. DAPI staining is shown as blue in merged images. Scale bar, 10 μm. (C and D) EGFP-FLAG-CD59 (C94S) and mRFP-KDEL constructs were transiently transfected into CANX&CALR-DKO cells carrying CANX mutants defective in N-glycan binding activity (Y164A, K166A, M188A or E216A) or defective in ERp57 binding (W342A&D343A, D343A&E351A or D347A&E351A). Confocal images were obtained at 3 days after transfection. Scale bar, 10 μm. (E) The GPI attachment signal of CD59 (C94S) was replaced with the transmembrane sequence of CD46 to generate a transmembrane form of misfolded CD59 (C94S) (EGFP-FLAG-CD59 (C94S) TM) and was co-transfected with mRFP-KDEL into WT and CANX&CALR-DKO cells. The localization of EGFP-FLAG-CD59 (C94S) TM was imaged by confocal microscopy. Scale bar, 10 μm.

**Figure 5-supplement.**
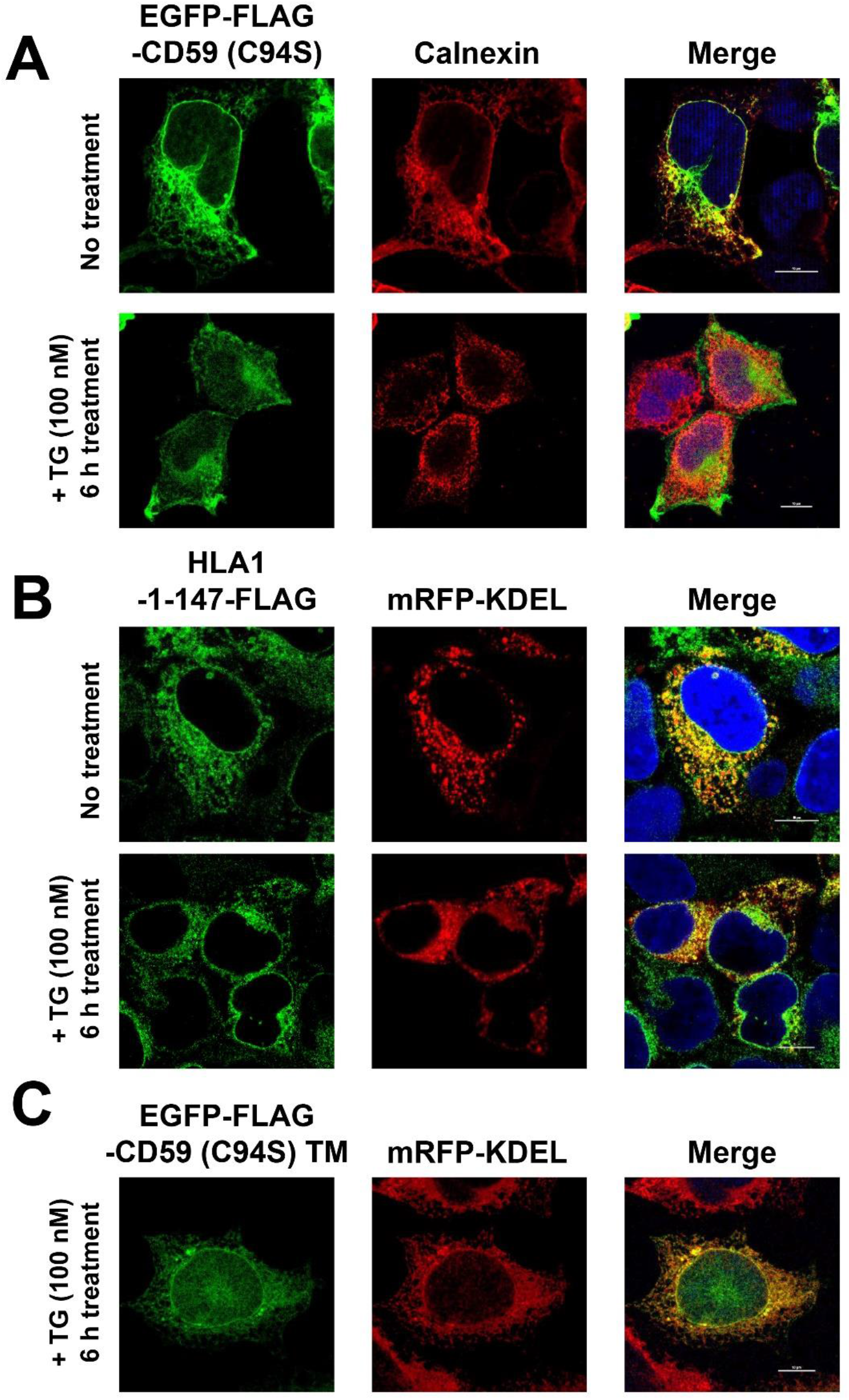
Protein localization change from the ER to the plasma membrane is specific for GPI-APs. (A) Localization of endogenous calnexin under acute ER stress. HEK293 cells transfected with EGFP-FLAG-CD59 (C94S) were incubated with or without 100 nM TG for 6 h. Cells were then fixed, stained with an anti-calnexin antibody followed by an anti-mouse-Alexa Fluor 555, and detected by confocal microscopy. Scale bar, 10 μm. (B and C) Localization of HLA1-1-147-FLAG (B) and EGFP-FLAG-CD59 (C94S) TM (C) under acute ER stress. HLA1-1-147-FLAG or EGFP-FLAG-CD59 (C94S) TM and mRFP-KDEL constructs were transfected into WT cells. Cells were then incubated with or without 100 nM TG for 6 h 3 days after transfection. HLA1-1-147-FLAG was stained with anti-FLAG antibody follow by anti-mouse-Alexa Fluor 488, and detected by confocal microscopy. EGFP-FLAG-CD59 (C94S) TM, and mRFP-KDEL were imaged by confocal microscopy. Scale bar, 10 μm.

**Figure 6-supplement.**
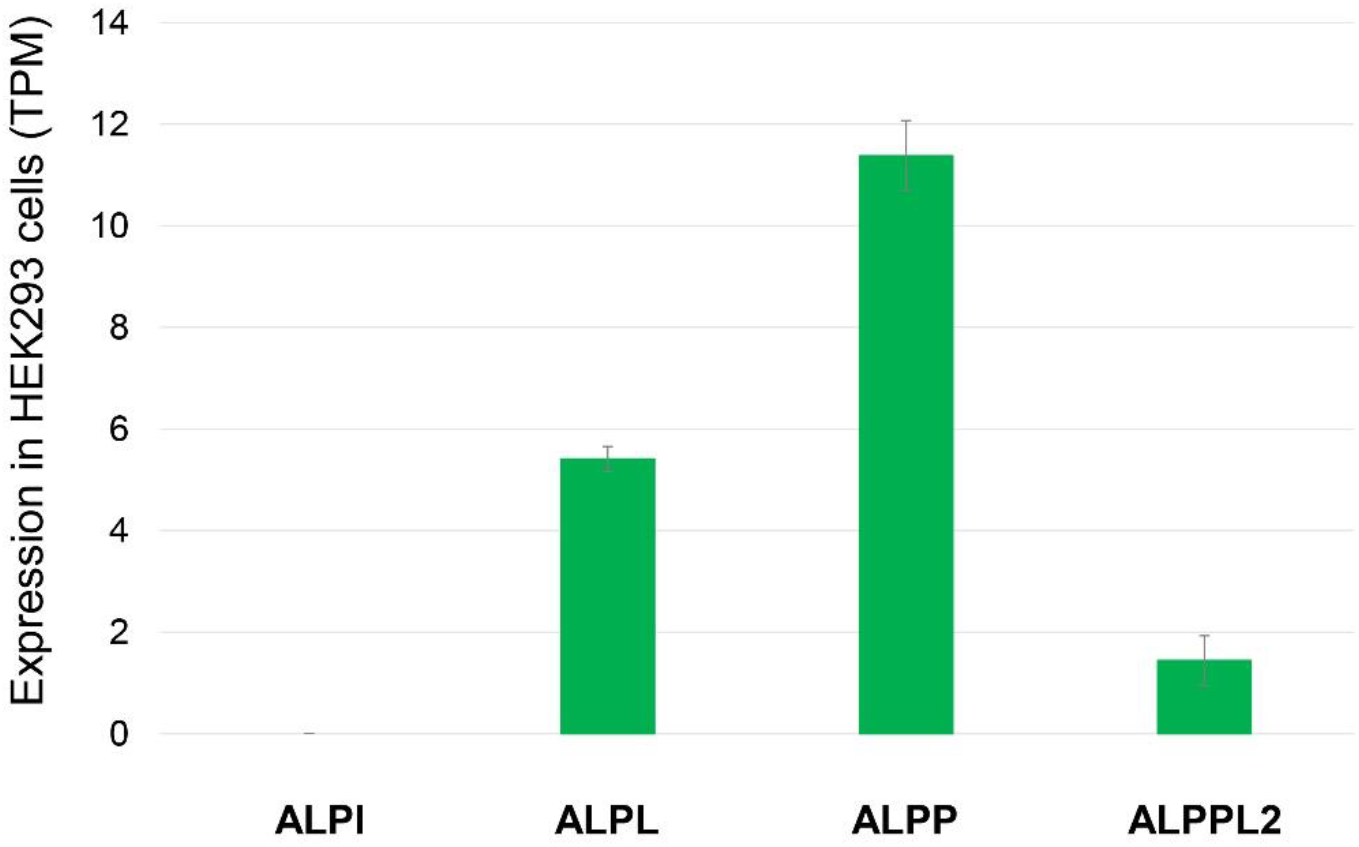
Expression of genes encoding alkaline phosphatases in HEK293 cells. The RNA-seq data of HEK293 cells (Huang et al., manuscript in preparation) were used to assess gene expression. Transcription per million (TPM) values are presented as the means ± SD of three independent measurements. ALPI, intestinal-type alkaline phosphatase; ALPL, tissue-nonspecific alkaline phosphatase; ALPP, placental type alkaline phosphatase; ALPPL2, germ cell type alkaline phosphatase.

